# Mitochondrial stress in GABAergic neurons non-cell autonomously regulates organismal health and aging

**DOI:** 10.1101/2024.03.20.585932

**Authors:** Laxmi Rathor, Shayla Curry, Youngyong Park, Taylor McElroy, Briana Robles, Yi Sheng, Wei-Wen Chen, Kisuk Min, Rui Xiao, Myon Hee Lee, Sung Min Han

## Abstract

Mitochondrial stress within the nervous system can trigger non-cell autonomous responses in peripheral tissues. However, the specific neurons involved and their impact on organismal aging and health have remained incompletely understood. Here, we demonstrate that mitochondrial stress in γ-aminobutyric acid-producing (GABAergic) neurons in *Caenorhabditis elegans* (*C. elegans*) is sufficient to significantly alter organismal lifespan, stress tolerance, and reproductive capabilities. This mitochondrial stress also leads to significant changes in mitochondrial mass, energy production, and levels of reactive oxygen species (ROS). DAF-16/FoxO activity is enhanced by GABAergic neuronal mitochondrial stress and mediates the induction of these non-cell-autonomous effects. Moreover, our findings indicate that GABA signaling operates within the same pathway as mitochondrial stress in GABAergic neurons, resulting in non-cell-autonomous alterations in organismal stress tolerance and longevity. In summary, these data suggest the crucial role of GABAergic neurons in detecting mitochondrial stress and orchestrating non-cell-autonomous changes throughout the organism.

## Introduction

Mitochondria are essential organelles involved in various cellular functions, including energy production, calcium regulation, cell signaling, and apoptosis. Their reciprocal relationship with aging is widely acknowledged, wherein mitochondrial dysfunction impacts organismal longevity and health, and aging affects mitochondrial homeostasis across variable model animals ^1, 2^. The degree of mitochondrial dysfunction determines the effect on an organismal lifespan. Mild mitochondrial disruption has been shown to increase lifespan in model animals such as *C. elegans*, flies, and mice ^3–10^. For instance, the lifespan of *C. elegans* is extended by inhibiting mitochondria-related genes through genetic and RNA interference (RNAi) knockdown, including *isp-1* (an iron-sulfur subunit of complex III of the mitochondrial electron transport chain (ETC)) ^6^, *spg-7* (a mitochondrial quality control m-AAA protease) ^11–13^, and *clk-1* (a hydroxylase involved in the biosynthesis of ubiquinone) ^5, 14^. Like *C. elegans clk-1* mutant, mice with the *mclk1* mutation, which exhibits normal growth and fertility, also exhibits an increase in lifespan ^15^.

The mechanisms underlying longevity enhancements through mitochondrial perturbations bring forth the mitohormesis theory that enhanced stress response pathways contribute to longer lifespans ^4, 16–21^. It has been demonstrated that some mitochondrial stress requires the mitochondrial unfolded protein response (mitoUPR) pathway to trigger lifespan extension ^4, 16–19^. The forkhead transcription factor DAF-16/FoxO has also been reported as an additional mediator responsible for extending the lifespan of *C. elegans* ETC mutants ^20, 21^. However, certain mitochondrial perturbations and mutations in ETC genes can independently extend lifespan without involving *atfs-1*, a key regulator of mitoUPR ^17, 22^. Similarly, it has been documented that DAF-16/FoxO is not indispensable in mediating the lifespan extension in some ETC mutants ^6^, suggesting a complex regulation of organismal aging in response to mitochondrial dysfunction through multiple pathways.

Aging is also accompanied by various changes in mitochondria, including the decreased activity of mitochondrial enzymes, reduced mitochondrial oxygen consumption, increased ROS production, decreased mitochondrial biogenesis, and increased mutations in mitochondrial DNA ^23–27^. Interestingly, different types and levels of mitochondrial DNA mutations accumulate in various tissues during aging in humans and model animals ^28, 29^. Even within a single tissue, such as the brain and muscle, variations in mitochondrial DNA mutations have been reported ^30–32^. As a result, there is growing interest in understanding how organisms respond to mitochondrial stress in specific tissues and cells.

Recent studies have elucidated the phenomenon wherein perturbations in mitochondrial function within a specific tissue can elicit lifespan extension and health improvement ^16, 33, 34^. Remarkably, the nervous system exhibits a particular role in detecting its intrinsic mitochondrial stress and extending organismal lifespan ^16, 33–36^. However, it remains unclear which neuronal subtype is responsible for extending organismal longevity in response to their mitochondrial stress and what mechanisms underlie the non-cell-autonomous changes in this regard.

γ-aminobutyric acid (GABA) is a widely utilized neurotransmitter found in both vertebrate and invertebrate nervous systems. In vertebrates, a substantial proportion, estimated to be between 30% and 40%, of central nervous system synapses rely on GABA transmission ^37^. GABA levels tend to gradually decrease as organisms age, and disruptions in GABA neurotransmission have been associated with various neurological disorders and age-related cognitive decline ^38, 39^. In *C. elegans*, 26 neurons have been anatomically and functionally characterized as GABAergic neurons ^40, 41^. Interestingly, recent studies in *C. elegans* indicate that the GABA signaling pathway plays an important role in regulating the lifespan and overall health of the organism, mitochondrial unfolded protein response, and proteostasis in the post-synaptic muscle tissue ^42–45^.

In this study, we utilized C. elegans as a model organism to explore the role of GABAergic neurons in regulating organismal health and aging in response to mitochondrial perturbations, as well as the mechanisms underlying these effects. Our findings indicate that mitochondrial stress in GABAergic neurons can influence the activity of DAF-16/FoxO in peripheral tissues through GABA signaling. This, in turn, leads to non-cell-autonomous alterations in the mitochondria activity and organismal healthspan, reproductive capacity, and lifespan.

## Results

### Mitochondrial perturbations in GABAergic neurons are sufficient to prolong the organismal lifespan

To explore the influence of mitochondrial dysfunction in GABAergic neurons on lifespan and healthspan, we inhibited the functions of *isp-1* and *spg-7*, well-documented genes that extend organismal lifespan when their normal functions are disrupted ^11–13, 33, 46, 47^. In line with previous studies, systemic RNA interference (hereinafter referred to as sRNAi) targeting *isp-1* and *spg-7* significantly extended the lifespan of wild-type N2 animals when compared to control animals subjected to empty vector (EV) control RNAi (Figures 1A and 1B; Table S1). Tissue-specific RNAi in GABAergic neurons (hereinafter referred to as gRNAi) was accomplished by employing a previously well-established transgenic animal ^48–50^. In this model, the function of *rde-1*, a member of the PIWI/STING/Argonaute protein family, was specifically restored in GABAergic neurons by introducing a transgene that expresses RDE-1 under the control of the GABAergic neuron-specific promoter derived from the *unc-47* gene (Pgaba) (Figure 1C) ^48, 51–53^. Additionally, they expressed *sid-1*, a gene responsible for dsRNA transport. When gRNAi against *isp-1* was induced from the L1 stage, it resulted in a median lifespan extension of 55.5% compared to the control (Figures 1D and Table S1). *spg-7(gRNAi)* animals also showed an extended median lifespan of 33.3% (Figure 1E and Table S1).

**Figure 1.**
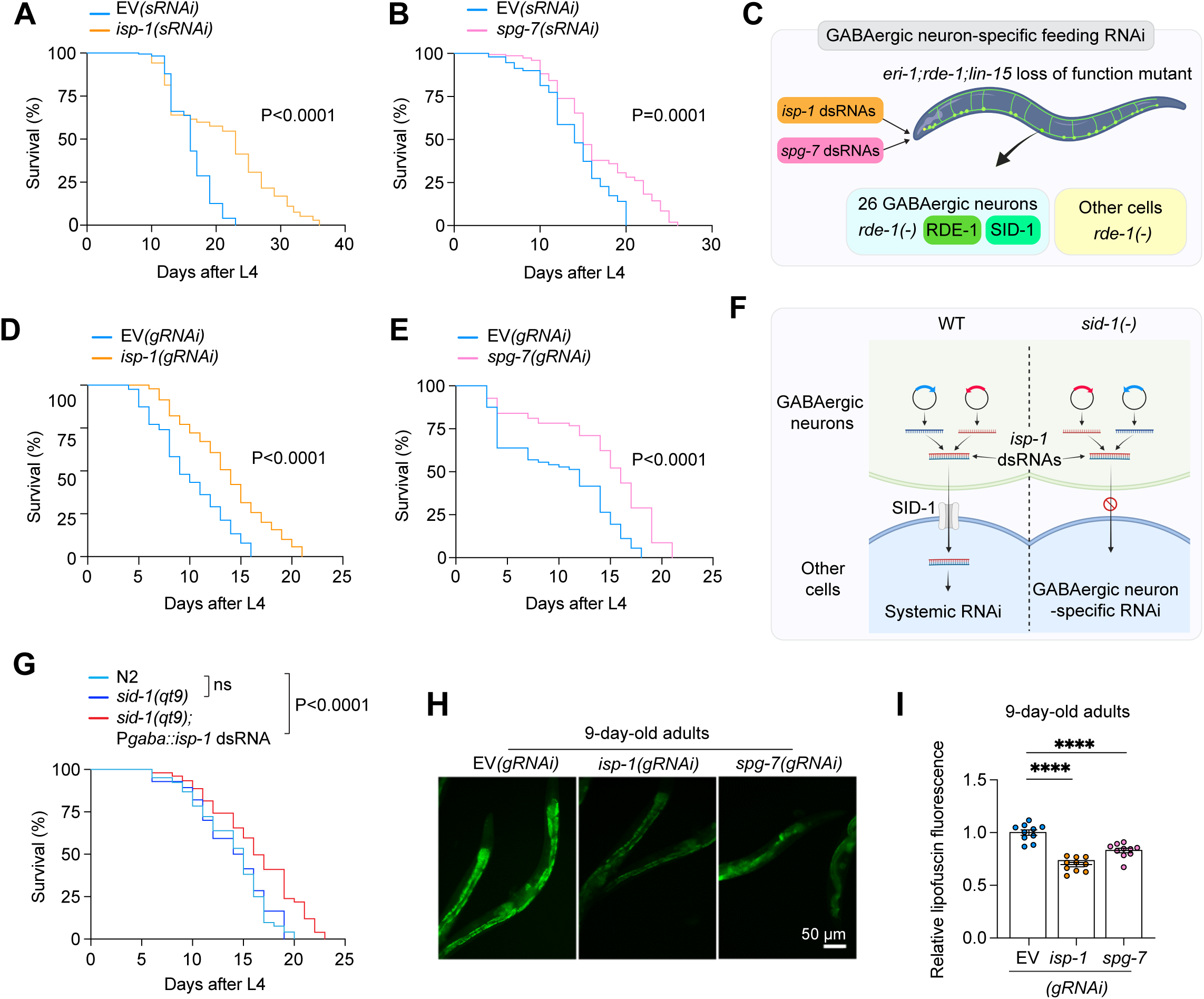
Mitochondrial stress in GABAergic neurons is sufficient to prolong the organismal lifespan. (A and B) Lifespan analysis of systemic *isp-1(sRNAi)* and *spg-7(sRNAi)* animals. (C) GABA-neuron-specific RNAi system. The gRNAi system is based on the *rde-1* mosaic mutant background, which carries a loss-of-function mutation in *rde-1*, thereby preventing RNAi ^48, 50, 51, 53, 56, 131^. This system also involves *eri-1* and *lin-15* loss-of-function mutant backgrounds, resulting in hyperactivated RNAi sensitivity in all neurons ^48, 132^. Restricted feeding RNAi to GABAergic neurons is achieved by exclusive RDE-1 expression in GABAergic neurons, along with the SID-1 dsRNA channel protein ^48^. (D and E) Lifespan analysis of *isp-1(gRNAi)* and *spg-7(gRNAi)* animals. (F) GABAergic neuron-*specific isp-1* RNAi through *in vivo* transcription of sense and antisense RNAs in GABAergic neurons under the *sid-1(eq9)* null mutant background. (G) Lifespan analysis of *sid-1(qt9)* and *sid-1(qt9)*+P*gaba::isp-1* dsRNA animals. The significance of the lifespan curves (A, B, D, E, and G) was assessed using a Log-rank (Mantel-Cox) test. (H and I) The levels of lipofuscin in *isp-1(gRNAi)* and *spg-7(gRNAi)* animals at the 9-day-old adult stage were compared to those in control animals. (H) Representative images of lipofuscin fluorescence in the whole body of animals under the indicated conditions. Bar, 50 µm. (I) Quantification of lipofuscin fluorescence intensity. Each dot represents a single animal. ****P < 0.0001; one-way ANOVA test. Data are expressed as means ± SEM. Created with https://www.biorender.com/.

To exclude the possibility of any remaining systemic RNAi effects in the *rde-1* mutant background, we carried out another tissue-specific RNAi strategy ^54^. We expressed *isp-1* double-stranded RNAs (dsRNAs) in GABAergic neurons using the Pgaba promoter (hereinafter referred to as P*gaba::isp-1* dsRNA) in the *sid-1(qt9)* null mutant background, which lacks intercellular dsRNA transport, preventing sRNAi (Figure 1F) ^55, 56^. The expression of *isp-1* dsRNAs in GABAergic neurons within the wild-type N2 animal background proved effective in upregulating the expression of P*hsp-6*::GFP, which serves as a fluorescent reporter for mitoUPR activity, specifically in the intestine, suggesting that this *isp-1* dsRNA expression effectively induced mitochondrial defects (Figure S1A and S1B) ^4, 34, 57^. Notably, we observed that P*gaba::isp-1* dsRNA expression in *sid-1* null mutants also significantly extended organismal lifespan (Figures 1G and Table S1) ^58–60^. Additionally, in 9-day-old adult *isp-1(gRNAi)* and *spg-7(gRNAi)* animals, we observed reduced lipofuscin fluorescence in the intestine, an aging hallmark that typically increases progressively over time (Figures 1H and 1I) ^61^. These findings collectively suggest that mitochondrial perturbation within GABAergic neurons is sufficient to prolong the organismal lifespan and attenuate the aging process.

### Mitochondrial stress in GABAergic neurons increases the stress tolerance of the organism

Next, we sought to determine whether mitochondrial stress in GABAergic neurons could also alter the parameters of a healthspan. We tested thermal and oxidative stresses during aging, which are healthspan parameters closely linked to longevity across species ^62–66^ (Figure 2A). In mid-age adults (3-day-old adult) groups, gRNAi against *isp-1* and *spg-7* increased survivability against paraquat exposure compared to controls (Figure 2B). This improvement was also observed in older adult groups (Figures 2C and 2D). Moreover, *isp-1(gRNAi)* and *spg-7(gRNAi)* animals displayed a significant increase in survival when exposed to thermal stress at 35 °C (Figures 2E-2G). To validate the efficacy of RNAi, we maximized the RNAi effect by culturing animals under the feeding RNAi condition against *spg-7* for three generations (Figure S2A) and observed consistently improved survival rates in response to both oxidative stress (Figures S2B and S2C) and thermal stress (Figures S3D and S3E). In addition, *sid-1* null mutants expressing P*gaba::isp-1* dsRNA expression also exhibited enhanced tolerance to thermal and paraquat stresses (Figures 2H and 2I). These findings suggest that mitochondrial stress in GABAergic neurons can enhance healthspan parameters.

**Figure 2.**
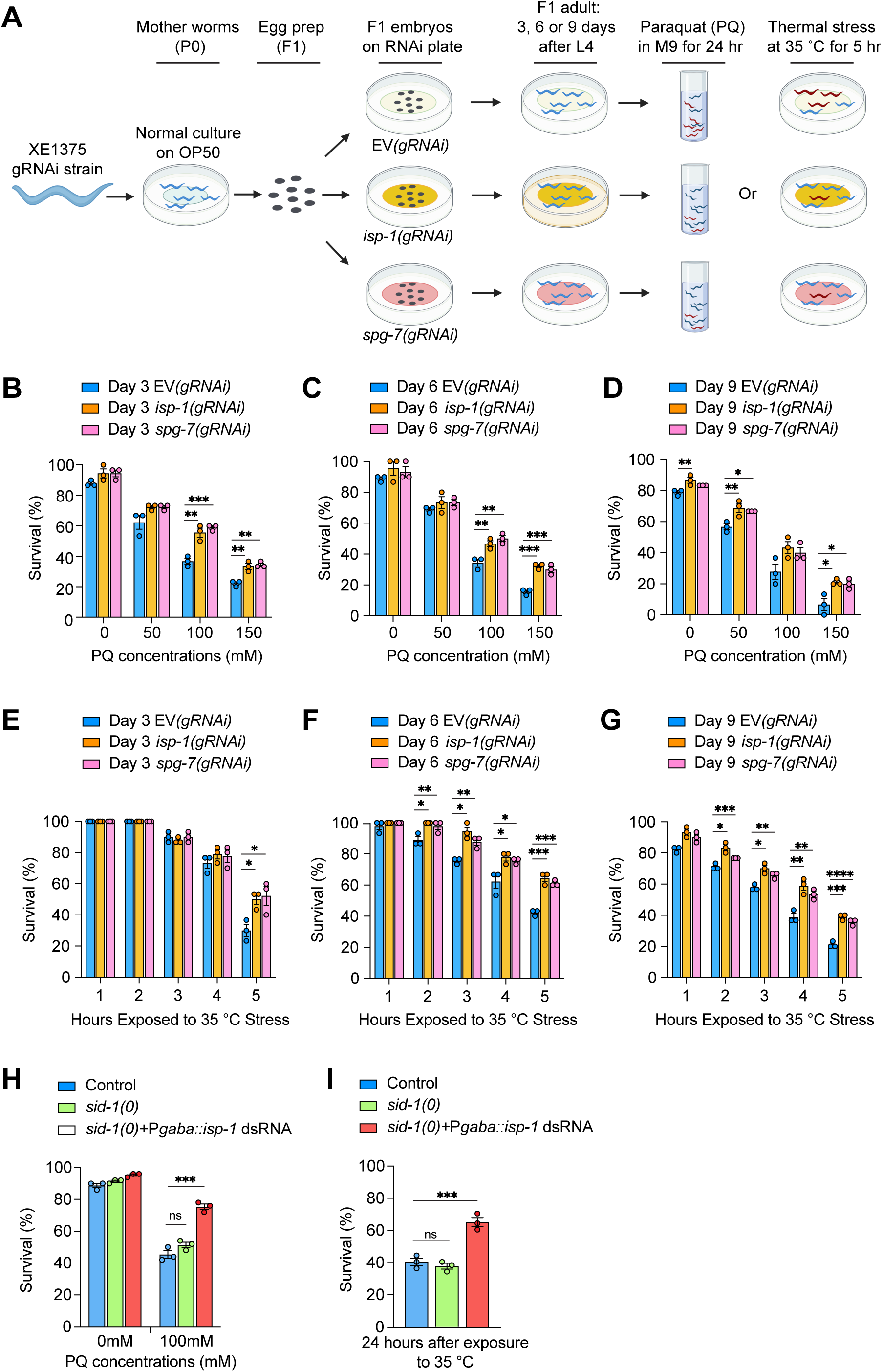
Mitochondrial stress in GABAergic neurons enhances stress resistance. (A) A diagram illustrating the paraquat and thermal stress experimental procedures is provided. (B-D) The survival rate of animals cultured in indicated concentrations of paraquat for 24 hours was measured at 3, 6, and 9-day-old adult stages. (E-G) The survival rate after incubation at 35°C for 5 hours was measured in animals at 3, 6, and 9-day-old adult stages. At least 30 animals were tested for each set. Statistical significance is indicated as follows: (H-I) Survival rates of animals at the 3-day-old adult stage after 100 mM paraquat treatment for 24 hours (H) and 24 hours after incubation at 35°C for 5 hours (I) were measured. *P < 0.05, **P < 0.005, ***P < 0.0005, ****P < 0.0001; one-way ANOVA test. Data are expressed as means ± SEM.

### GABAergic neuronal mitochondrial stress alters reproduction

While depletion of some mitochondrial ETC in the entire nervous system of *C. elegans* and flies increases lifespan in a non-cell-autonomous manner, it does not consistently affect fertility ^8, 16^. Therefore, we sought to determine if mitochondrial dysfunction in GABAergic neurons similarly has no impact on reproduction. *C. elegans* exists as a hermaphrodite and produces a limited number of sperm in the L4 stage. In the adult stage, it undergoes oocyte development and self-fertilizes to produce embryos (Figure 3A) ^67, 68^. While *isp-1* mutant has a prolonged lifespan, it has been shown to have a reduced brood size ^69^. Interestingly, *isp-1* gRNAi was sufficient to reduce the number of fertile animals (Figure 3B). The total number of embryos produced by fertile *isp-1(gRNAi)* animals was also significantly decreased (P < 0.0001) (Figure 3C). When we assessed reproductive activity daily, fertile *isp-1(gRNAi)* animals displayed a reproductive period similar to that of the control group. However, brood sizes from the 2-day-old stage to the 4-day-old stage were significantly reduced (P<0.005 for the 2 and 3-day-old stages and P<0.0001 for the 4-day-old stage) (Figure 3D). Intrigued by these results, we evaluated the impact of *isp-1(gRNAi)* on the three critical stages of germline development including mitotic germ cell proliferation, meiotic germ cell apoptosis, and oogenesis (Figure 3A). Although the length of the mitotic area was not significantly affected by *isp-1(gRNAi)* (Figures 3E and S3A), the total number of mitotic germ cells was decreased during the early reproductive periods (1 to 3 days after the L4 stage) (Figure 3F). The number of apoptotic germ cells in the meiotic gonadal loop region, undergoing germline apoptosis, was not different from that in control animals at the 2-day-old adult stage, but it was substantially decreased at the 3-day-old stage (Figure 3G) ^70^. Thus, an increased loss of meiotic germ cells was not likely the primary cause of the reduced brood size. Interestingly, 2-day-old adult *isp-1(gRNAi)* animals displayed large DNA aggregates in the proximal gonad, a characteristic feature of an endomitotic oocyte phenotype (Figures 3H and 3I)^71^. Once embryos were produced, most of them were hatched in *isp-1(gRNAi)* animals (Figure S3B). Our results indicated that disruption of the mitochondrial ETC in GABAergic neurons negatively affected germline development and reproductive ability.

**Figure 3.**
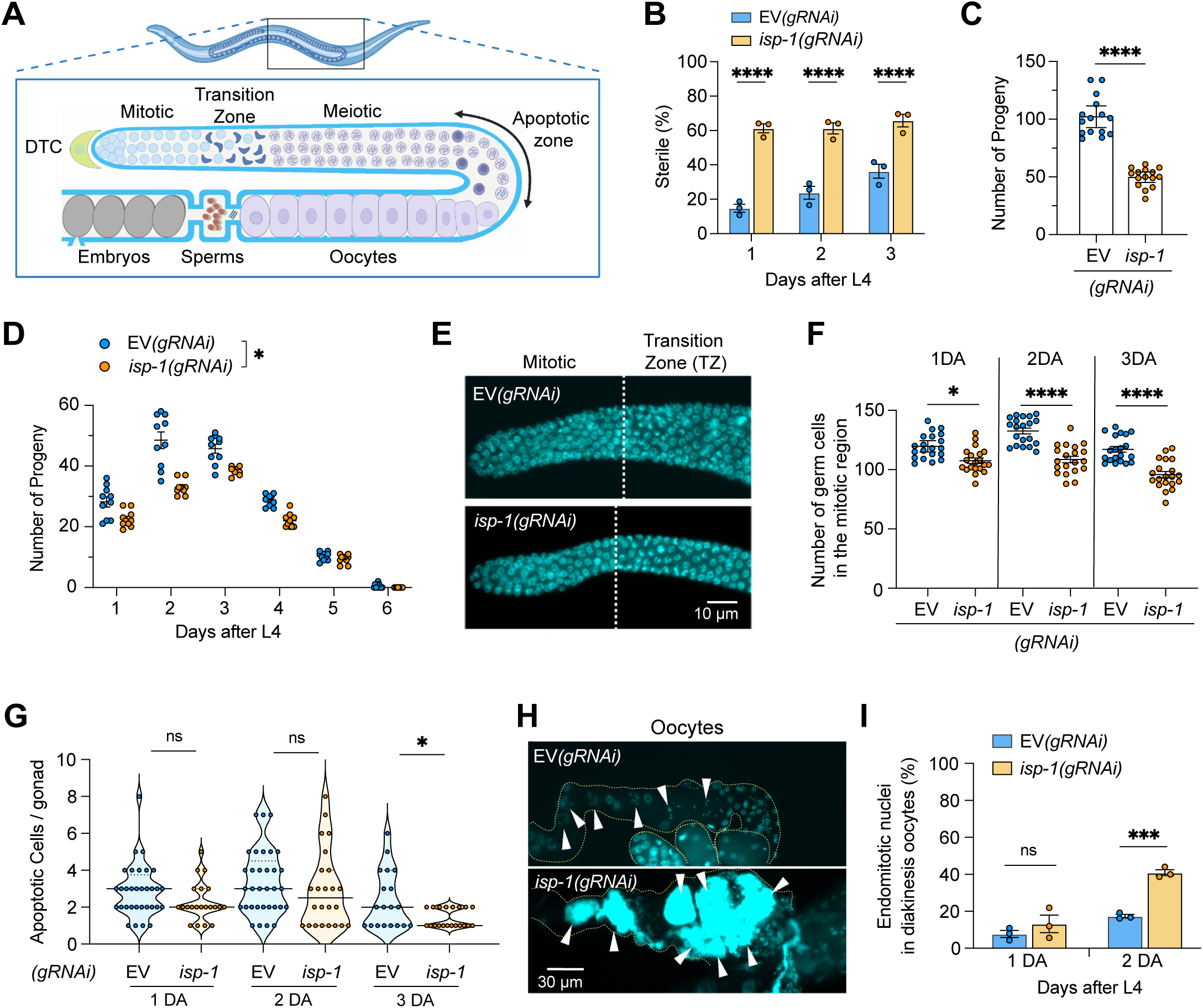
Reproductive effects of mitochondrial perturbation in GABAergic neurons. (A) A diagram of the *C. elegans* hermaphrodite reproductive system was presented, showing the distal gonad with the mitotic area for germ cell proliferation, the meiotic germ area for meiosis, and an area undergoing germ cell apoptosis, the proximal area with oogenesis, the spermatheca for sperm storage, and the uterus for fertilized egg storage. (B) Sterility in *isp-1(gRNAi)* animals compared to control animals. (C) The total number of embryos in fertile *isp-1(gRNAi)* and *spg-7(gRNAi)* animals. (D) The daily brood size in fertile *isp-1(gRNAi)* and *spg-7(gRNAi)* animals. (E) Representative images of dissected distal gonad arms stained with DAPI from 2-day-old *isp-1(gRNAi)* and control animals. The dashed lines indicate the endpoint of the mitotic area. (F) Quantitative analysis of germ cell numbers in the mitotic regions of *isp-1(gRNAi)* and control animals. (G) A violin plot is presented, displaying the median (solid line) and quartiles (dashed lines), illustrating the apoptotic germ cells in the loop regions of *isp-1(gRNAi)* and control animals. (H) Representative images of dissected proximal gonad stained with DAPI from 2-day-old *isp-1(gRNAi)* and control animals. The arrowheads indicate stained oocyte chromosomes. (I) Quantitative analysis of endomitotic nuclei in diakinesis oocytes in the proximal gonad of *isp-1(gRNAi)* and control animals. Statistical significance is indicated as follows: *P < 0.05, ***P < 0.0005, ****P < 0.0001; two-tailed Mann–Whitney test. Each dot represents an individual worm (C and D) and gonad arm (F, G, and I). Additionally, each dot indicates an individual group (B and I), animal (C and D), and gonad (F and G). Data are expressed as means ± SEM except (G). Created with https://www.biorender.com/.

### Mitochondrial stress in GABAergic neurons enhances mitochondrial function in the peripheral tissues

Accumulated evidence indicates that aging is linked to a decline in mitochondrial function and biogenesis ^72–75^. Additionally, experimentally increasing mitochondrial membrane potential and biogenesis is associated with lifespan extension ^75, 76^. Therefore, we hypothesized that the mitochondrial stress specific to GABAergic neurons could impact the function and homeostasis of mitochondria in other peripheral tissues. At the 2-day-old stage, staining animals with the mitochondrial membrane potential-dependent MitoTracker Red CMXRos dye revealed a higher mitochondrial membrane potential in both the whole body (Figures 4A and 4B) and the intestine (Figure 4C) of *isp-1(gRNAi)* and *spg-7(gRNAi)* animals compared to that in the control group ^77,78^. Additionally, whole-body extracts from *isp-1(gRNAi)* animals showed a marked increase in ATP levels (Figure 4D). *spg-7(gRNAi)* animals showed somewhat higher ATP levels compared to the control group, but it was not significant (Figure 4D). Next, we stained *isp-1(gRNAi)* and *spg-7(gRNAi)* animals using the MitoTracker FM Green dye that accumulates in mitochondria in a membrane potential-independent manner, indicating the mass of mitochondria ^79^. We observed that both *isp-1(gRNAi)* and *spg-7(gRNAi)* animals exhibited higher MitoTracker FM Green fluorescence in the whole body than the control animals, suggesting an increase in mitochondrial mass (Figures 4E and 4F). In agreement with these results, the copy number of mitochondrial DNA was increased in *isp-1(gRNAi)* and *spg-7(gRNAi)* animals compared to control animals (Figure 4G). The expression of mitochondrial DNA polymerase gamma *polg-1* was also significantly upregulated in *isp-1(gRNAi)* animals (Figure 4H) ^35, 80, 81^. Notably, staining *isp-1(gRNAi)* and *spg-7(gRNAi)* animals with the ROS indicator DCF-DA showed lower ROS levels compared to the control animals (Figures 4I-4K) ^82^. Altogether, these results suggest that disturbances in mitochondrial homeostasis within GABAergic neurons could systemically increase overall mitochondrial membrane potential, ATP levels, and mitochondrial mass while decreasing ROS levels.

**Figure 4.**
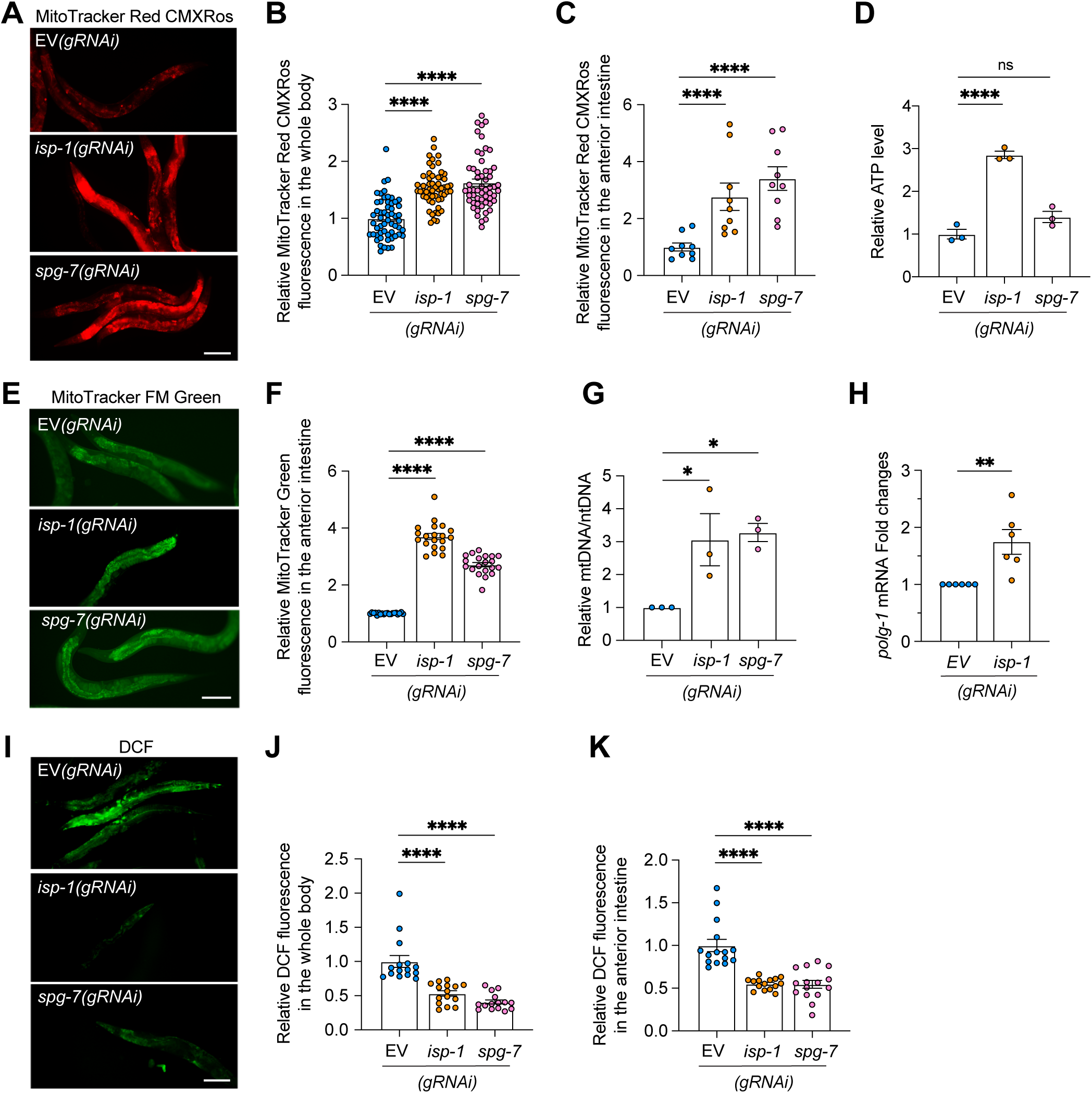
GABAergic neuronal mitochondrial stress alters the mitochondrial homeostasis in peripheral tissues of animals. (A) Representative images of mitochondrial membrane potential in *isp-1(gRNAi)* and *spg-7(gRNAi)* animals after MitoTracker CMXRos dye staining at 3-day-old adults. (B and C) Quantitative analysis of MitoTracker CMXRos fluorescent intensity measured by microscopy with a 100x magnification in the indicated animals at 3-day-old adult stage (B) and in the anterior intestine region assessed by microscopy with a 400x magnification (C). (D) The ATP bioluminescence assay measured ATP levels in the whole-body extracts of *isp-1(gRNAi)* and *spg-7(gRNAi)* animals. (E) Representative image of MitoTracker FM Green fluorescence in animals at 3-day-old adults to visualize mitochondrial mass. (F) Quantitative MitoTracker FM Green fluorescence intensity in *isp-1(gRNAi)* and *spg-7(gRNAi)* animals (G) Relative mtDNA copy number analyzed by the qPCR method in *isp-1(gRNAi)* and *spg-7(gRNAi)* animals. (H) Relative transcript levels of mitochondrial DNA polymerase gamma *polg-1* gene expression measured by qPCR in *isp-1(gRNAi)* animals at the 2-day-old adult stage. (I) Representative images of *isp-1(gRNAi)* and *spg-7(gRNAi)* animals stained with H2DCF-DA dye to measure ROS levels in the whole body. (J) Quantitative analysis of H2DCF fluorescent intensity monitored by a spectrophotometer in *isp-1(gRNAi)* and *spg-7(gRNAi)* animals at 3-day-old adults. (K) Quantitative fluorescence intensity in the anterior intestine of animals after H2DCF-DA staining with a 400x magnification. Statistical significance is indicated as follows: *P < 0.05, **P < 0.005, ***P < 0.0005, ****P < 0.0001; one-way ANOVA test for (B, C, D, F, G, J, and K) and two-tailed student’s t-test for (H). Each dot indicates an individual worm (B, C, F, J, and K). Data are expressed as means ± SEM. Bars, 50 µm.

### GABAergic neuronal Mitochondrial Stress Enhances the DAF-16/FoxO Pathway

Despite the increased total mitochondrial mass and activity, the decreased ROS levels in *isp-1(gRNAi)* and *spg-7(gRNAi)* animals suggest the possibility that their ability to mitigate oxidative stress has improved. Therefore, we analyzed changes in stress response regulators associated with mitochondria and ROS, including DAF-16/FoxO, mitoUPR, and SKN-1/Nrf ^21, 57, 83–85^. It has been demonstrated that mutations in *isp-1* and RNAi targeting *isp-1* (*isp-1* sRNAi) result in enhanced expression of reporter genes associated with the mitoUPR pathway, such as *hsp-6* or *hsp-60* ^4, 34, 57^. In line with this, when the expression of P*gaba*::*isp-1* dsRNA was induced in N2 wild-type worms, leading to systemic *isp-1* RNAi, a significant increase in P*hsp-6*::GFP expression was observed (Figures S1A and S1B). Additionally, we observed that sRNAi against *isp-1* effectively triggered the expression of P*hsp-6*::GFP, and this induction was dependent on *atfs-1*, a crucial mediator of the mitoUPR pathway ^86^ (Figure S4A). These findings indicate the efficiency of our RNAi feeding conditions. However, *isp-1* gRNAi did not significantly elevate the mRNA levels of *hsp-60* and *hsp-6,* as evaluated by quantitative reverse transcription PCR (RT-qPCR) analysis (Figure 5A). The mRNA level of *gst-4*, a downstream target of the SKN-1/Nrf pathway, was also not significantly increased by *isp-1(gRNAi)* (Figure 5A) ^87, 88^. In contrast, the mRNA levels of three DAF-16/FoxO downstream genes, including *sod-3*, *hsp-16.2*, and *dlk-1*, were substantially increased by *isp-1(gRNAi)* (Figure 5A) ^89–91^. Additionally, *sid-1(eq9)* null mutants expressing *isp-1* dsRNA in GABAergic neurons also showed increased expression of DAF-16/FoxO target genes (Figure 5B). The intensity of a fluorescent reporter for the DAF-16/FoxO pathway (*muIs84* [P*sod-3::gfp*]) was also enhanced in this condition (Figures 5C and 5D). The elevated expression of *sod-3* and *dlk-1* induced by *isp-1(gRNAi)* was suppressed by the loss of DAF-16, suggesting that their expression is mediated by the DAF-16/FoxO pathway (Figure 5E).

**Figure 5.**
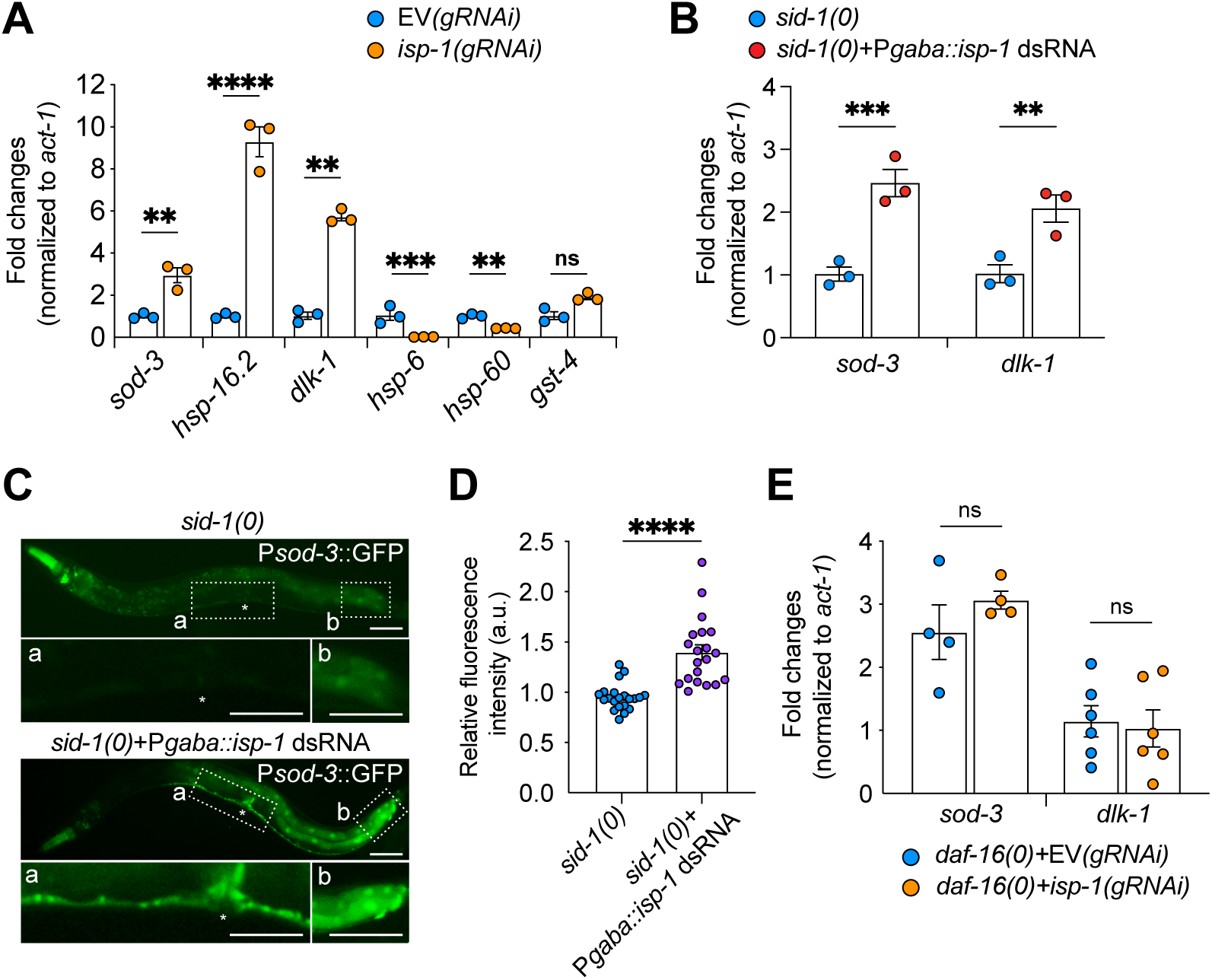
GABAergic neuronal mitochondria stress elevated the DAF-16/FoxO pathway. (A) RT-qPCR analysis to assess the relative transcript levels of mitoUPR, SKN-1/Nrf, and DAF-16/FoxO signaling target genes in *isp-1(gRNAi)* animals compared to control EV*(gRNAi)* animals. The expression levels of each gene were normalized to the reference gene *act-2* which encodes actin. (B) RT-qPCR analysis was conducted to assess the relative transcript levels of *sod-3* in *sid-1(qt9)* null mutants expressing *isp-1* dsRNA in GABAergic neurons. (C and D) Representative images and the quantification of P*sod-3::*GFP expression, a reporter for DAF-16/FoxO activity, in *sid-1(qt9)* control animals and *sid-1(qt9)+* P*gaba::isp-1* dsRNA animals. Each dot indicates an individual worm. (E) RT-qPCR analysis was performed to assess the relative transcript levels of *sod-3* and *dlk-1* in *daf-16(mgDf47)* null mutants, both with and without *isp-1* gRNAi treatment. Statistical significance is indicated as follows: **P < 0.005, ***P < 0.0005, ****P < 0.0001; two-tailed student’s t-test (B); two-tailed Mann–Whitney test (D); two-way ANOVA (A and E). Data are expressed as means ± SEM.

### The non-cell-autonomous effects of GABAergic neuronal mitochondrial defects require DAF-16/FoxO

Further investigations were conducted to evaluate the functional involvement of DAF-16/FoxO in the non-cell-autonomous effects of mitochondrial stress in GABAergic neurons. It was found that sRNAi against *isp-1* required DAF-16/FoxO function to extend lifespan, as previously reported (Figure S4B and Table S1) ^20, 21^. DAF-16/FoxO was also necessary for the enhanced tolerance against paraquat in *isp-1(sRNAi)* worms (Figure S4C). Notably, the lifespan extension by *isp-1* gRNAi was suppressed in *daf-16(mgDf47)* null mutants (Figure 6A). Additionally, DAF-16/FoxO was required for the increased stress tolerance against thermal (Figure 6B) and paraquat (Figure 6C) stresses in *isp-1(gRNAi)* animals. DAF-16 loss also suppressed the upregulation of mitochondrial membrane potential (Figures 6D and 6E) and mitochondrial mass (Figures 6F and 6G) induced by *isp-1* gRNAi. Moreover, gRNAi against *isp-1* in the *daf-16(mgDf47)* null mutant background failed to increase mitochondrial DNA copy number (Figure 6H) and *polg-1* mRNA levels (Figure 6I). Finally, the *daf-16* null mutation suppressed the abnormalities in daily reproductive ability (Figure 6J) and total brood size (Figure 6K) caused by *isp-1* gRNAi. Interestingly, the double RNAi knockdown of *isp-1* and *daf-16* in GABAergic neurons (*isp-1*+*daf-16* gRNAi) also resulted in a normal lifespan, to a level similar to that displayed in EV*(gRNAi)* and *daf-16(gRNAi)* conditions, suggesting that DAF-16 function in GABAergic neurons is required to mediate the lifespan extension (Figure S4D and Table S1). These findings collectively suggest that DAF-16/FoxO plays a critical role in mediating non-cell autonomous changes resulting from GABAergic neuronal mitochondrial stress, influencing organismal lifespan, stress tolerance, mitochondrial homeostasis, and reproductive capacity.

**Figure 6.**
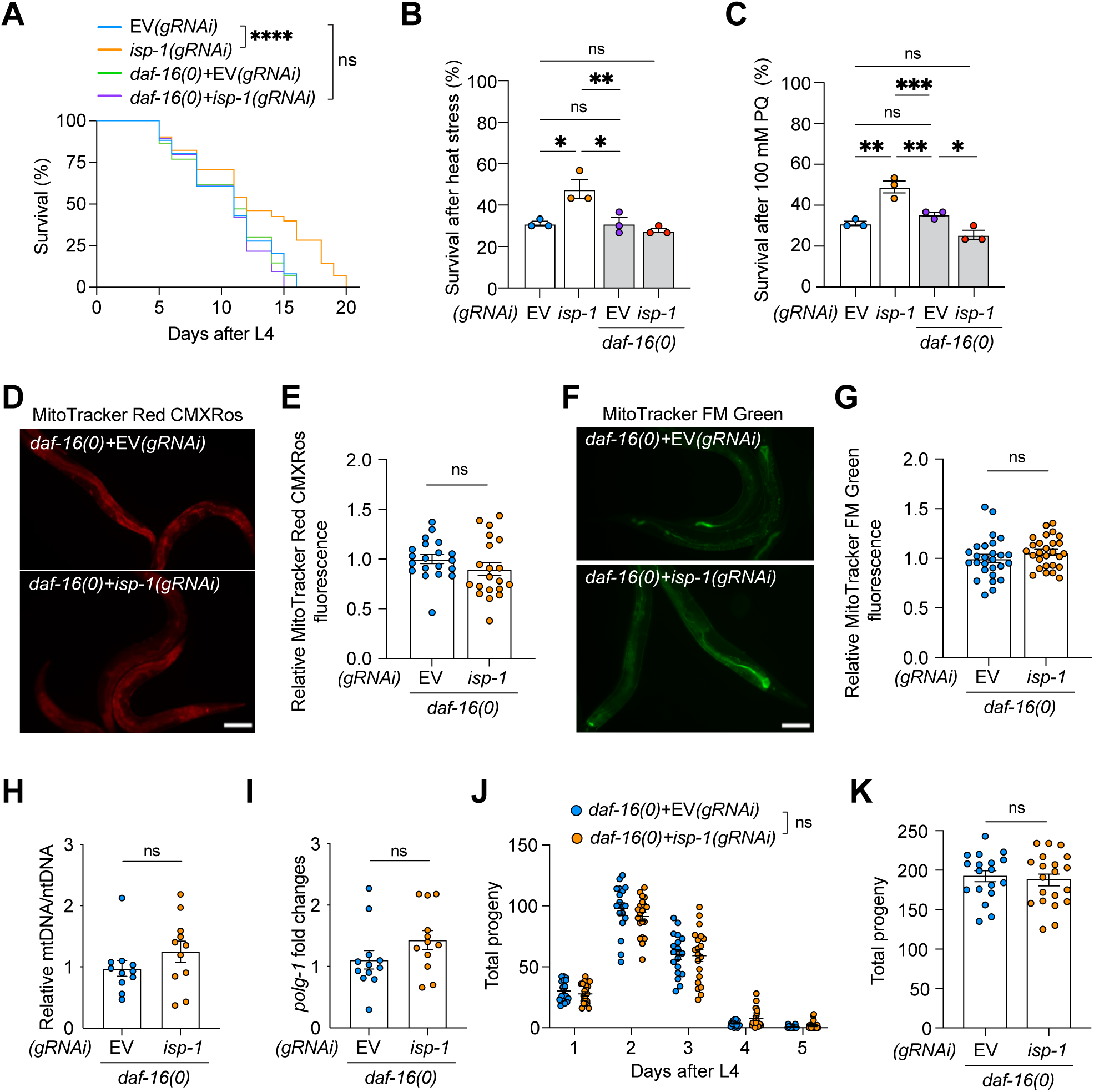
GABAergic neuronal mitochondria stress requires DAF-16/FoxO to trigger the non-cell-autonomous effects. (A) Lifespan analysis to evaluate the role of *daf-16* in the extended lifespan induced by P*gaba::isp-1* dsRNA. The significance of the lifespan curves was assessed using a Log-rank (Mantel-Cox) test. (B and C) Stress tolerance assays. In a *daf-16(mgDf47)* mutant background, P*gaba::isp-1* dsRNA expression failed to increase resistance to heat stress (B) or paraquat stress (C) compared to control EV gRNAi. (D and E) Representative images (D) and quantification (E) of mitochondrial membrane potential assessed by MitoTracker Red CMXRos dye staining at 3-day-old adults. (F and G) Representative images (F) and quantification (G) of MitoTracker FM Green fluorescent staining at 3-day-old adults to monitor mitochondria mass. (H and I) Relative mtDNA copy number (H) and *polg-1* gene expression level (I) were analyzed by qPCR in EV(gRNAi) and *isp-1(gRNAi)* under *daf-16(mgDf47)* null mutant background. (J) Comparison of the mean number of eggs laid each day between *daf-16(mgDf47)*+*isp-1(gRNAi)* and control *daf-16(mgDf47)*; EV*(gRNAi)* animals. (K) The total number of embryos in fertile *daf-16(mgDf47)*+*isp-1(gRNAi)* and control *daf-16(mgDf47)*; EV*(gRNAi)* animals. Each dot represents an individual animal (E, G, J, and K). Statistical significance is indicated as follows: *P < 0.05, **P < 0.005, ***P < 0.0005, ****P < 0.0001; a one-way ANOVA test for B and C, and a two-tailed Mann–Whitney test for E, G, H, I, J, and K. Data are expressed as means ± SEM. Bars, 50 µm.

### Mitochondrial stress in GABAergic neurons causes non-cell-autonomous changes by acting on the same mechanisms as GABA signaling

Recent studies in *C. elegans* have revealed that GABA signaling plays a role not only in the regulation of GABAergic neuronal function but also in governing organismal longevity. Specifically, loss of *unc-25*, which encodes glutamic acid decarboxylase, has been shown to prolong lifespan and enhance stress tolerance ^42, 43^. Hence, we conducted epistatic tests to investigate the possible interaction between the loss of GABA signaling and *isp-1* knockdown conditions in GABAergic neurons and their impact on organismal health and longevity. We found that targeting *isp-1* specifically in GABAergic neurons by expressing P*gaba::isp-1* dsRNAs in *sid-1(qt9)* mutants did not lead to a further extension of lifespan in *unc-25* null mutants (Figures 7A and S5A; Table S1). Additionally, GABAergic neuronal mitochondrial stress did not lead to an additive increase in stress tolerance beyond the levels observed in *unc-25(e156); sid-1(qt9)* mutants (Figures 7B and 7C). The enhanced GFP expression driven by the *sod-3* promoter due to *isp-1* dsRNA expression in GABAergic neurons was not further increased in the *unc-25* null mutant condition (Figure 7D).

**Figure 7.**
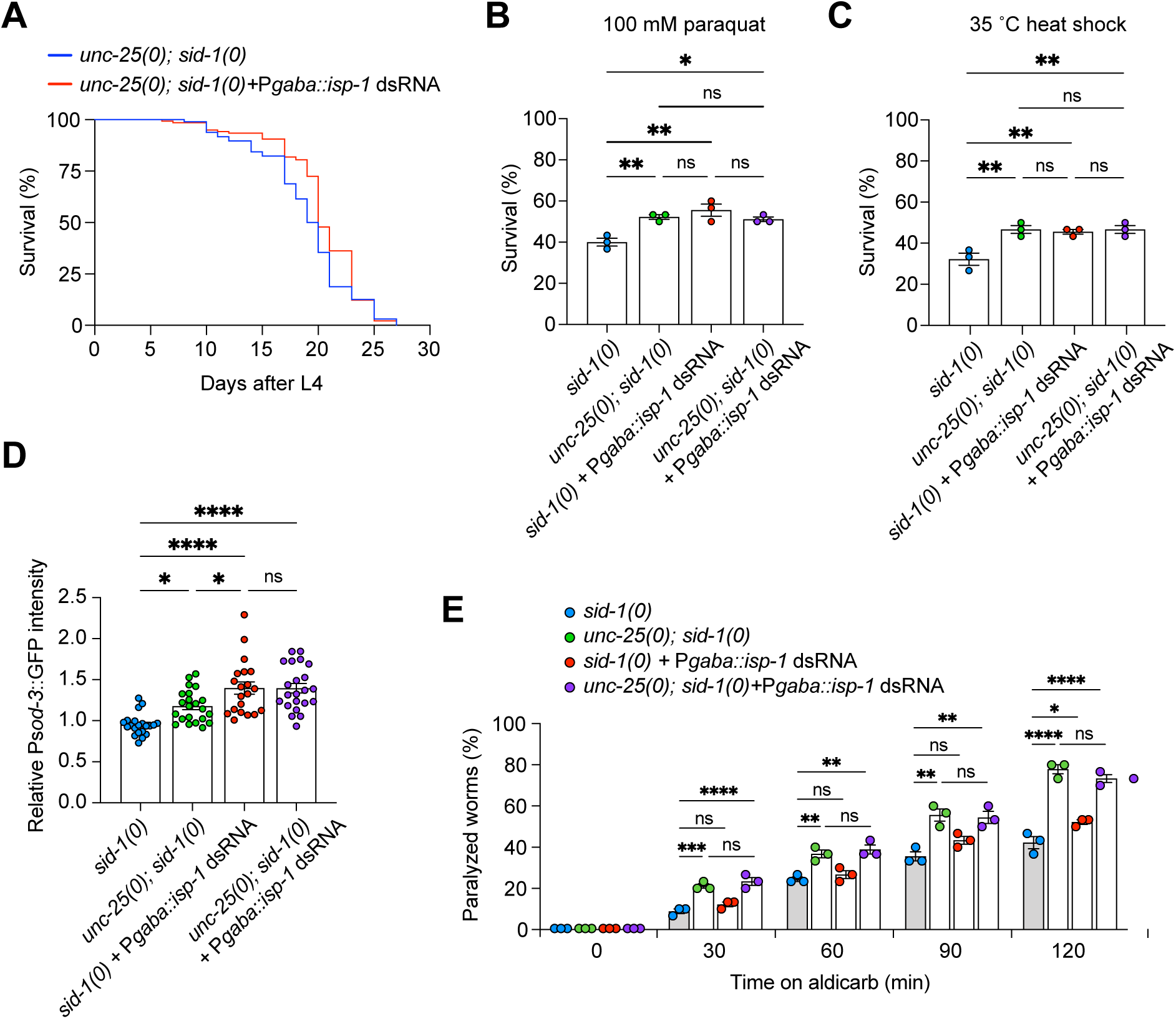
Non-cell autonomous effects of GABAergic neuronal mitochondrial stress were not additively enhanced by the loss of GABA signaling. (A) Lifespan analysis was conducted on animals with indicated genetic backgrounds to investigate the relationship between GABA signaling and mitochondrial stress in GABAergic neurons. (B and C) Paraquat and thermal stress assays were performed on *unc-25* null mutants with and without GABAergic neuronal mitochondrial stress induced by P*gaba::isp-1* dsRNA expression to elucidate their relationship. (D) Assessment of the fluorescence intensity of the P*sod-3*::GFP reporter, indicative of DAF-16/FoxO activity, in the specified mutants. (E) Evaluation of aldicarb sensitivity in response to the expression of *isp-1* dsRNAs in GABAergic neurons. Each dot represents an individual group in B, D, and E and individual animal in D. *P < 0.05, **P < 0.005, ***P < 0.005, ****P < 0.001; Log-rank (Mantel-Cox) test for A; one-way ANOVA test for B, C, D, and E. Data were expressed as means ± SEM.

Next, we assess if GABAergic neuronal mitochondrial stress affects GABA function by testing the sensitivity of animals to aldicarb, an acetylcholinesterase inhibitor that can cause post-synaptic receptor hyperstimulation and paralysis due to reduced acetylcholine breakdown ^92^. As previously reported, depletion of GABA in *unc-25* mutants increased the sensitivity to aldicarb ^92–94^ (Figures 7E). GABAergic neuronal expression of *isp-1* dsRNA also significantly elevated the sensitivity to aldicarb, particularly when animals were exposed to it for a prolonged period (120 min). The expression of P*gaba*::*isp-1* dsRNA did not further increase the hypersensitivity to aldicarb in *unc-25* mutants, indicating that they function in the same pathway. Together, these findings suggest that diminished GABA signaling and mitochondrial stress within GABAergic neurons trigger alterations in organismal lifespan and healthspan through a common pathway.

### Neuropeptide signaling in GABAergic neurons regulates organismal aging and health without additive effects with mitochondrial stress in GABAergic neurons

Next, we tested the potential roles of neuropeptide signaling in the non-cell autonomous effects of GABA neuronal mitochondrial stress ^33^. We measured the lifespan of mutants with mutations in *unc-31*, required for dense-core vesicle exocytosis ^95^. In agreement with the previous report, *unc-31(e928)* mutants exhibited prolonged lifespan (Figure 8A and Table S1). There is no additive lifespan increase by P*gaba*::*isp-1* dsRNA expression in the *unc-31* mutant background, suggesting the potential involvement of neuropeptide signaling in regulating lifespan (Figure 8A and Table S1). GABAergic neurons have been suggested to express several neuropeptides, including *flp-10, flp-11, flp-13*, and *flp-22* ^96, 97^. In line with previous studies, *flp-13* was expressed in a subset of *C. elegans* GABAergic motor neurons (Figure S6A) ^97, 98^. We found that *flp-13*+EV gRNAi extended the lifespan of animals compared to control animals, indicating its non-cell autonomous function in GABAergic neurons in regulating organismal lifespan (Figure 8B); Table S1). Double gRNAi against *flp-13* and *isp-1* did not further increase lifespan compared to *flp-13*+EV gRNAi alone, suggesting that depletion of FLP-13 and mitochondrial disruption in GABA neurons could extend lifespan through a common mechanism (Figure 8B and Table S1). While gRNAi against *flp-13*+EV did not affect the tolerance of animals against heat stress, it improved stress tolerance against paraquat to a level comparable to that observed in animals treated with *flp-13*+*isp-1* double gRNAi (Figures 8C and 8D). Maximal treatment of *flp-13* gRNAi, without mixing with EV bacteria, did not significantly increase the heat tolerance compared to the control group, suggesting that the lack of impact of *flp-13*+EV gRNAi on heat stress is unlikely to be attributed to a diluted RNAi efficiency due to the double RNAi method (Figure S6B). Additionally, maximal treatment of *flp-13* gRNAi still showed enhanced paraquat tolerance and lifespan (Figures S6C and S6D; Table S1). Collectively, these findings indicate a novel role for the neuropeptide FLP-13 in regulating organismal aging and health. This regulation appears to be mediated by a common mechanism associated with non-cell autonomous effects resulting from GABA neuronal mitochondrial dysfunction. The selective response of *flp-13* mutants to specific stressors implies the existence of additional mechanisms that respond to mitochondrial stress within GABAergic neurons.

**Figure 8.**
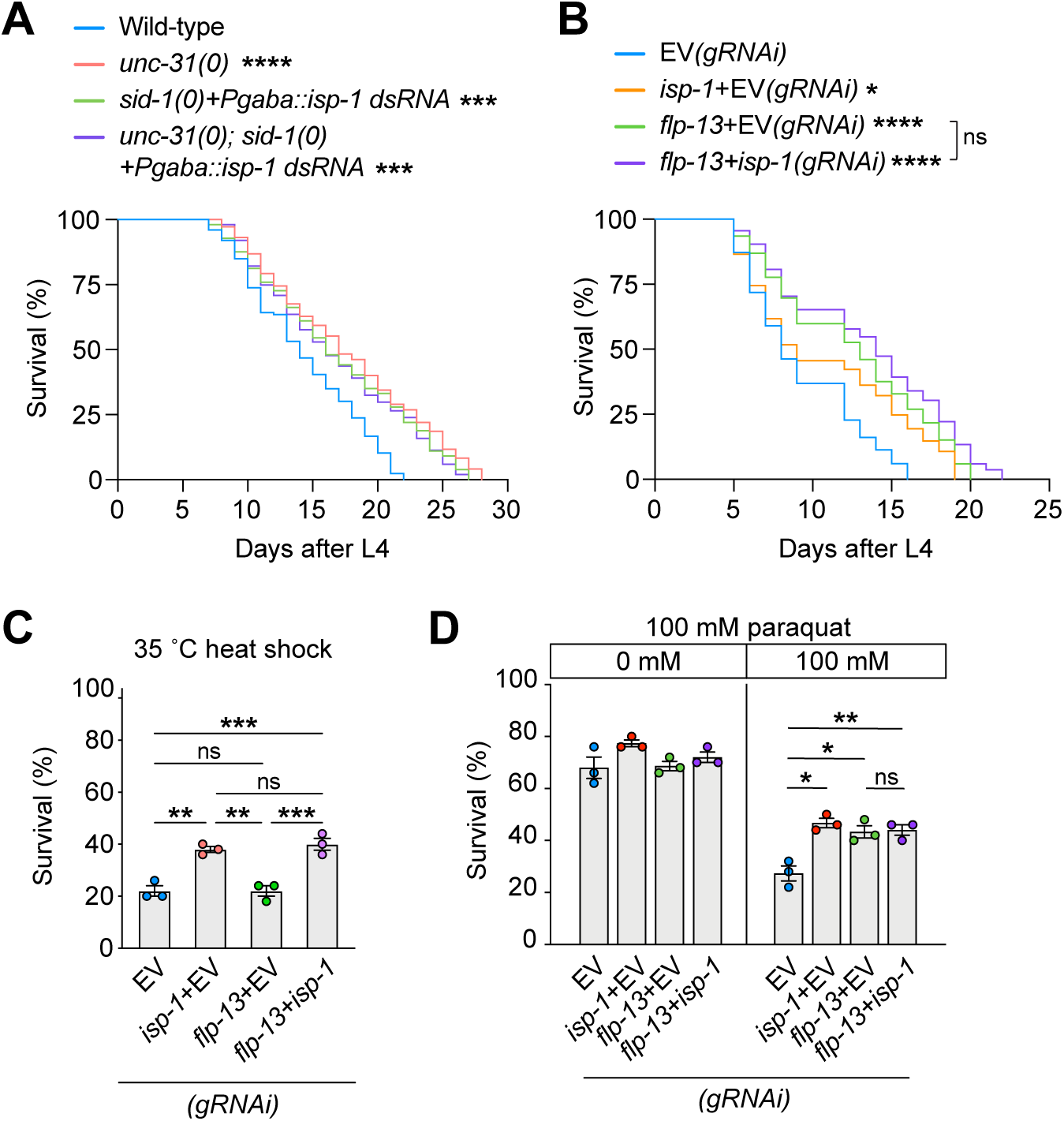
FLP-13 neuropeptides modulate organismal lifespan and stress tolerance, without expanding the non-cell autonomous effects of GABAergic neuronal mitochondrial stress. (A and B) Lifespan assays were conducted to evaluate the role of *unc-31* (A) and *flp-13* in regulating organismal longevity. (C and D) Quantitative analysis of the thermal (C) and paraquat (D) stress resistance assays was performed after double gRNAi, induced by feeding worms a 50:50 mixture of two bacterial strains expressing either EV, *flp-13*, or *isp-1* dsRNAs. *P < 0.05, **P < 0.005, ***P < 0.005, ****P < 0.001; Log-rank (Mantel-Cox) test for A and B; one-way ANOVA test for H and I. Data were expressed as means ± SEM.

## Discussion

Our findings suggest that GABAergic neurons play a critical role in sensing mitochondrial stress and regulating longevity. Additionally, disrupting the mitochondria in GABAergic neurons alone was sufficient to induce alterations in organismal health and reproduction. Enhanced DAF-16/FoxO activity is necessary for mediating the non-cell autonomous changes observed in animals. Furthermore, we identified GABA signaling and one neuropeptide signaling as a factor associated with the non-cell autonomous effects caused by mitochondrial stress in GABAergic neurons.

Previous studies have demonstrated that mitochondrial stress in all neurons is sufficient to extend organismal lifespan. However, it remains incompletely understood which neurons respond to their mitochondrial stress to mediate this non-cell autonomous effects on organismal lifespan. Sha and colleagues have tested 18 neurons out of the total 302 in *C. elegans* regarding their potential to activate non-cell-autonomous effects and demonstrated that a specific subset, consisting of ASK, AWA, AWC, and AIA neurons, exhibits the capacity to induce non-cell-autonomous activation of the mitoUPR pathway in response to mitochondrial stress their mitochondria perturbation ^33^. Interestingly, these conditions do not alter organismal lifespan, suggesting an unlink between mitoUPR and lifespan in certain conditions ^33^.

The specific role of GABAergic neurons in non-cell autonomous regulation of aging and health in response to mitochondrial dysfunction has not been reported. This could be because these GABAergic neurons have not been tested or that previous studies primarily focused on neurons that non-cell autonomously affect the mitoUPR pathway in the peripheral tissues ^16, 33–36^. However, a recent study reveals that induced ROS production in GABAergic neurons can trigger mitoUPR induction in other peripheral tissues through activating UNC-49/GABA_A_ receptor signaling, suggesting a central role of GABAergic neurons in regulating organismal stress response ^45^. In our study, we prioritized monitoring changes in the lifespan rather than detecting mitoUPR reporter activity and found that inducing mitochondrial stress, specifically in GABAergic neurons, was sufficient to extend the lifespan of the animal. We employed two independent RNAi strategies to knock down two genes related to mitochondria. The tissue-specific RNAi approach, utilizing the *rde-1(ne219)* null mutant background and exclusively restoring *rde-1* in target tissues, has been successfully utilized in multiple studies ^48, 50, 51, 53^. To eliminate the possibility of incomplete gRNAi, we also conducted *isp-1* knockdown in GABAergic neurons by expressing *isp-1* dsRNAs in *sid-1* null mutant backgrounds, and we consistently observed an extension in lifespan in both RNAi strategies. *isp-1* mutants and P*gaba::isp-1* dsRNA expression in wildtype animals robustly increase mitoUPR. In contrast, our qPCR results showed that there was no increase in mitoUPR reporter expression in *isp-1(gRNAi)* animals ^4^. Additionally, the *isp-1(gRNAi)* animals did not fully recapitulate the phenotypes of *isp-1* mutants, including normal ATP level, increased ROS level, and decreased mitochondrial membrane potential ^99–102^, supporting that our two *isp-1* knockdown conditions did not occur systemically other than in GABAergic neurons.

Longevity is associated with healthspan, but not in all cases ^103^. Our results show that mitochondrial stress in GABAergic neurons not only prolonged lifespan but also improved stress tolerance, a typical healthspan parameter. Notably, we also found enhancements in mitochondria activity and ATP levels. These improvements in mitochondria could be, in part, a result of an increase in mitochondria population evidenced by increases in mitochondrial DNA copy number and mass. Consistently this notion, we also found increased mRNA levels of *polg-1*, encoding the mitochondrial DNA polymerase that is responsible for the replication of the mitochondrial genome, suggesting a potential increase in mitochondria biogenesis by GABAergic neuronal mitochondrial stress ^35, 104^. The concept of aging in model organisms has long been linked to a decrease in mitochondrial function and biogenesis ^1^. Similarly, in humans, mitochondrial function declines with age ^2, 105^. Enhancing mitochondrial function has been proposed as an intervention strategy against aging. In mice, the brain-specific overexpression of *Sirt1*, a pivotal regulator of mitochondrial biogenesis through PGC-1a, not only extended lifespan but also induced non-cell-autonomous effects in skeletal muscle including improved mitochondria homeostasis ^75^. A recent study indicates that experimentally rejuvenated mitochondrial membrane potential is sufficient to extend *C. elegans* lifespan ^76^. Additionally, while *isp-1* mutants have been shown to have elevated levels of ROS, a central factor in aging, our findings suggest that knockdown of *isp-*1 in GABAergic neurons leads to a reduction in ROS levels, despite an increased mitochondrial population and heightened activity ^6, 102, 106^. Therefore, both enhanced mitochondrial homeostasis and reduced ROS levels could contribute to the non-cell-autonomous enhancement in aging and health.

The reduced ROS also proposed the potential involvement of the stress response pathway against ROS. Strikingly, our qPCR results showed no evidence supporting the activation of mitoUPR by GABAergic-neuronal mitochondrial stress ^16, 33–36^. It has been shown that *afts-1* acts cell-autonomously in certain neurons to mediate non-cell autonomous effects of mitochondrial stress in specific neurons ^33^. Thus, it is still possible that mitoUPR activation in GABAergic neurons plays a role in mediating lifespan extension and improving stress resistance. Recent studies have demonstrated that experimentally enhanced ROS production in GABAergic neurons activates the mitoUPR and this requires UNC-49 GABA_A_ receptor ^45^. However, *unc-49* mutants exhibit a normal lifespan ^42, 43^. Therefore, the extended lifespan resulting from mitochondrial defects in GABAergic neurons is not primarily mediated by increased ROS production and non-cell-autonomous activation of mitoUPR.

Our results suggested that GABAergic neuronal mitochondrial stress requires DAF-16/FoxO. We found that GABA-neuronal mitochondrial stress promoted the expression of *sod-3, hsp-16.2,* and *dlk-1*, which are regulated by DAF-16 ^89–91^. A recent study also reveals that ROS induction in GABAergic neurons results in non-cell autonomously increased expression of *sod-3* along with *hsp-6* ^45^. DAF-16 is another pathway suggested to be activated by mitochondrial dysfunction, contributing to the extended lifespan observed in *C. elegans* with whole-body mitochondrial dysfunction ^21^. However, another study reports that while DAF-16 is required for the longevity of mitochondrial mutants in certain conditions, it was not sufficient to fully account for the observed lifespan extension ^6^. We found that depletion of DAF-16 suppressed a series of non-cell-autonomous changes in lifespan, stress resistance, mitochondrial DNA (mtDNA) copy number, and *polg-1* expression. These results indicate the functional importance of DAF-16 in mediating the non-cell autonomous effects of mitochondrial disruption in GABAergic neurons.

Recent studies on *C. elegans* have reported that GABA signaling regulates lifespan and health ^42, 43^. Notably, GABA loss increases DAF-16/FoxO activity in the intestine, thereby increasing organismal longevity and health span ^42, 43^. Additionally, GABA signaling modulates protein homeostasis in *C. elegans* post-synaptic muscle cells ^44^. In contrast to the role of GABA signaling in *C. elegans* lifespan, knockdown of the *Drosophila* GABA_B_ receptor shorts lifespan ^107^. Nevertheless, these results indicate that GABA signals could have a conserved role in regulating organismal health and aging. Our aldicarb assay results suggest that GABAergic neuronal mitochondrial stress could interfere with GABAergic neuronal activity. It has been revealed that mitochondria abundantly accumulate at the presynapse and affect synaptic activity by regulating the supply of ATP and calcium homeostasis, which are required for proper neurotransmitter release and recycling ^108–113^. Our previous studies have demonstrated that over 75% of mitochondria localize at the presynapse in GABAergic motor neurons in *C. elegans* ^114^. A screen for essential genes required for GABAergic neuronal function, utilizing a GABAergic neuron-specific feeding RNAi in *C. elegans*, has identified several mitochondrial-related genes. Knockdown of these genes in GABAergic neurons triggers hypersensitivity against aldicarb ^48^. Therefore, GABAergic neuronal mitochondrial perturbation could mimic effects induced by a reduction in GABA signaling. This hypothesis is supported by our epistatic analysis results, suggesting that GABAergic neuronal mitochondrial stress and the loss of GABA signaling in *unc-25* mutants may share a common mechanism influencing longevity, stress tolerance, and *sod-3* expression. Understanding how mitochondrial stress can modulate downstream mediators such as DAF-16 through altering GABA singling and ultimately affecting longevity and health can be complex. Mitochondrial stress in GABAergic neurons may partially reduce GABA signaling to varying degrees rather than completely turning it off ^112, 115^. Moreover, GABA signaling differentially regulates lifespan and each healthspan parameter through three receptors and a combination of four downstream pathways ^43^. Thus, reduced GABA signaling caused by GABA mitochondrial stress could affect each downstream receptor pathway to varying degrees, depending on the ability of each receptor to respond to GABA ligands.

It has been well-documented that cell-autonomous responses to mitochondrial defects can vary depending on the stressor conditions, including disruptions in each ETC component and disruption modes, pathological conditions, and environmental stressors ^101, 116^. These variations highlight the complex responses to mitochondrial disturbances. Interestingly, the non-cell autonomous longevity changes also differ depending on the type of mitochondrial perturbation. It has been reported that inhibition of *spg-7* and *cco-1,* encoding a cytochrome c oxidase-1, in the entire nervous system prolongs lifespan ^16, 33^, but it was not suppressed by mitoUPR loss. Mitochondrial stress induced by pan-neuronal expression of polyglutamine repeats (polyQ40) and mutation in *ucp-4,* encoding a mitochondrial uncoupling protein-4, do not further prolong the lifespan than wild-type animals, but mitoUPR function is required to preserve normal lifespan ^33^. Also, the deletion of *spg-7* in AIY neurons is sufficient to induce mitoUPR in distal tissues but does not prolong lifespan ^7,32,35^. Additionally, pan-neuronal expression of KillerRed, which induces oxidative stress, resulted in a reduced lifespan, despite an enhanced mitoUPR pathway in remote tissues ^33^. In this study, we targeted to inhibition of *isp-1* and *spg-7* in GABAergic neurons and observed consistent changes in lifespan and stress resistance. Notably, *isp-1* gRNAi resulted in decreased reproductive ability. Previous studies report that depletion of CCO-1, a component of mitochondria ETC, in the nervous system results in mitoUPR activation in the peripheral tissues and prolongs lifespan without affecting brood size ^16, 33^. Similarly, in *Drosophila*, ETC reduction in the nervous system increases longevity but maintains normal fertility ^8^. However, it remains unclear whether other mitochondrial stressors in GABAergic neurons can produce similar non-cell-autonomous effects as seen with *isp-1* and *spg-7* knockdown, which could be one reason the role of GABAergic neurons in regulating the non-cell-autonomous effects of mitochondrial disturbance has not been elucidated previously.

Finally, studies have demonstrated that neuropeptide signaling mediates the non-cell autonomous regulation of mitoUPR and organismal aging in response to mitochondrial stress in specific neurons ^42, 117–122^. For instance, mitochondrial stress in ASK, AWA, AWC, and AIA neurons utilizes the FLP-2 neuropeptide to induce mitoUPR in peripheral tissues ^33^. These results suggest that there could be additional signaling molecules mediating the non-cell autonomous effects of GABAergic neuronal mitochondrial stress. GABAergic neurons have been found to express several neuropeptides, including *flp-10, flp-11, flp-13*, and *flp-22* ^96, 97^. Our finding indicates that gRNAi against *unc-31* increased lifespan resulting from depletion of dense core vesicle exocytosis was not further increased by GABA-neuronal mitochondrial stress, suggesting that neuropeptide signaling could be involved in non-autonomous changes caused by GABA-neuronal mitochondrial damage ^95^. FLP-13 loss increased lifespan and stress resistance, which is similar to the phenotype of animals with mitochondrial perturbations in GABAergic neurons. Animals with both conditions did show significant changes in stress resistance and lifespan compared to animals with either single condition, indicating that they work through the same mechanism to mediate non-cell autonomous effects. Therefore, it is possible that GABAergic neuronal mitochondrial stress reduces FLP-13 function, which inhibits lifespan and healthspan. Future studies are needed to elucidate the specific role of FLP-13 and other neuropeptides in the non-cell-autonomous effects of GABA-neuronal mitochondrial stress.

## Acknowledgment

We thank Drs. Shinichi Someya, Christiaan Leeuwenburgh, Stephanie Wohlgemuth, and Sejin Lee for their valuable discussions. Some strains were provided by the CGC, funded by the NIH Office of Research Infrastructure Programs (P40 OD010440). This research was conducted while SMH was a Hevolution/AFAR New Investigator Awardee in Aging Biology and Geroscience Research (AWD16056). SMH was also supported by the National Institute on Aging (NIA) of the National Institutes of Health (NIH) under award numbers R56AG066654 and R01AG081270, as well as the Glenn Foundation for Medical Research and the American Federation for Aging Research under award numbers AWD06577 and AWD11009 to SMH. Additionally, support was provided by NIA/NIH under award number AG060373-01 and the National Science Foundation under award number IOS (2132286) to MHL.

## Author Contributions

S.M.H. supervised the study. M.H.L. and S.M.H. conceived the study. L.R. and S.M.H. wrote the manuscript with input from all authors. L.R. and S.C. designed and performed experiments and analyzed and interpreted data. Y. P. and M.H.L designed germline experiments and interpreted data. B.R. and T.M. assisted with planning and performing the stress response and lifespan assays. Y.S. and R.X generated reagents and assisted with planning and interpreting data. K.M., W.C., and R.X. provided expertise and resources and reviewed the manuscript.

## Inclusion and diversity

We support inclusive, diverse, and equitable conduct of research.

## Data availability

All numerical data supporting the findings of this study are available in the supplemental information and from the corresponding author upon reasonable request.

## Material and Methods

### C. elegans strains

All *C. elegans* strains were maintained at 20 °C on nematode growth medium (NGM) plates seeded with the OP50 strain of *Escherichia coli* (*E. coli*) as described before ^123^. A list of strains used in this study is the following: N2 (Wild type), XE1375 (*wpIs36* I; *wpSi1* II; *eri-1(mg366)* IV; *rde-1(ne219)* V; *lin-15B(n744)* X), SJ4100 (*zcIs13* [*hsp-6p::gfp*]), CB156 (*unc-25(e156)* III), HC196 (*sid-1(qt9)* V), GR1352 (*daf-16(mgDf47)*I), TJ356 (*zIs356* IV [*daf-16p::daf-16a/b::gfp; rol-6(su1006)*]), CF1553 (muIs84 [*(pAD76) sod-3p::gfp + rol-6(su1006)*]), CL2166 (*dvIs19* III [*(pAF15)gst-4p::gfp::NLS*]). HAN260 (*daf-16(mgDf47) I*; wpIs36 I; *wpSi*1 II; *eri-1(mg366)* IV; *rde-1(ne219)* V; *lin-15B(n744)* X), HAN186 (*unc-25(e156); sid-1(qt9))*, HAN356 (*unc-25(e156); sid-1(qt9)*+ *sbsEx27* [*unc-47p::isp-1 sense+ unc-47p::isp-1 antisense*]), HAN357 (*sid-1(qt9)*+*sbsEx27* [*unc-47p::isp-1 sense+ unc-47p::isp-1 antisense*]).

### RNAi assay

RNAi was performed by the feeding method ^124, 125^. Briefly, freshly streaked single colonies of HT115(DE3) bacteria containing either empty L4440 vector (control) or *isp-1* and *spg-7* RNAi plasmid were grown overnight at 37°C in Luria broth (LB) medium supplemented with carbenicillin (25 μg/ml). HT115(DE3) bacterial feeding strains were obtained from the genome-wide library ^125^. PCR and sequencing were used to confirm that strains contained the correct clones. RNAi bacteria were seeded on NGM plates containing IPTG (1 mM) and cultured overnight before transferring animals. To prevent undesired non-specific effects, we did not use 5-fluoro-2′-deoxyuridine (FUdR) in seeded RNAi plates. Duplex RNAi was performed by mixing two HT115(DE3) bacterial strains, each containing the desired RNAi plasmid or empty vector in a 1:1 volume ratio. The efficiency of double RNAi, which involves feeding a 1:1 volume ratio mixture of bacteria strains targeting two different genes, was compared to that of single gene RNAi, which involves feeding a 1:1 volume ratio mixture of bacteria strains targeting one gene and an empty vector.

### Lifespan assay

Age-synchronized animals were prepared by egg prep from adult animals cultured on each RNAi or regular NGM plate at 20°C ^126^. The isolated embryos were transferred and allowed to hatch on RNAi NGM plates for the desired gene or regular NGM plates. Then, approximately 50 animals at the L4 stage were transferred to fresh RNAi plates. Every day, animals that failed to respond to gentle prodding with platinum wire were scored as dead. Lifespan data were statistically analyzed for significance by the log-rank test, comparing survival curves using GraphPad Prism software. Lifespan assays were performed at least in triplicate. To prevent undesired non-specific effects, we did not use 5-fluoro-2′-deoxyuridine (FUdR) in the seeded RNAi plates.

### Aldicarb sensitivity assay

Aldicarb sensitivity assay was performed as described previously ^92^. Briefly, approximately 40-50 mutant animals were grown on OP50-seeded NGM plates at 20 °C for 72 hours. Approximately 40 L4 stage mutant animals were then picked and placed on 35 mm OP50-seeded NGM plates containing 0.5 mM aldicarb (Sigma Aldrich, 33386: prepared aldicarb NGM plates 1 day before the assay) and scored for paralysis every 30 minutes over a 120-minute period. Animals were considered paralyzed when they failed to show any movements in response to touching at a fixed time. This assay was carried out in triplicate for each experimental condition.

### Reproductive assay

After pre-exposure to *spg-7* and *isp-1* RNAi from the L1 stage, age-synchronized XE1375 animals at the L4 stage (n=∼20-30) were individually transferred to fresh *spg-7* and *isp-1* RNAi NGM plates. Every day, we moved mother animals to new *spg-7* and *isp-1* RNAi NGM plates until egg-laying stopped. The embryos produced daily were counted, and the total number of produced embryos was used to calculate the brood size. The egg production period was used to calculate the reproductive span, and the number of embryos produced each day was used to analyze reproductive trends. The hatching rate was scored 24 hours after egg-laying. The brood size, reproductive span, hatching rate, and reproductive trend data were generated from independent experiments. At least three replicate experiments were performed for each assay.

### Stress assays

For the paraquat assay, 30-50 age-synchronized animals were pretreated to RNAi for each gene from the L1 stage. Then, 2-day-old adult animals were transferred to 96-well plates containing M9 with paraquat, a reactive oxygen species generator (Sigma-Aldrich, 36541-100MG), at a concentration of 0 mM, 50 mM, 100 mM, and 150 mM in a total volume of 100 µl per well. The survival of animals was evaluated after 24 hours, and animals that failed to respond to platinum wire touch were scored as dead. In the heat stress resistance assay, age-synchronized adult animals pretreated with *isp-1* or *spg-7* RNAi were exposed to thermal stress at 37°C for 5 hours. The survival after heat shock was recorded every hour for 5 hours by gently prodding with a platinum loop. Animals that failed to respond were scored as dead. All experiment was independently repeated at least three times.

### Staining assays for ROS level and mitochondrial homeostasis

We used the fluorescent probe H_2_DCF-DA dye (Invitrogen, D399) to detect ROS levels *in vivo*, as previously described ^82^. 2-day-old adult stage mutants or XE1375 animals pretreated with *spg-7*, *isp-1,* or control EV RNAi from the L1 stage were washed with M9 buffer to remove the bacteria. After washing 3 times, animals were collected in 300 µl of PTw buffer (1xPBS with 0.1% tween20). Then, around 30-35 animals were transferred to 96-well plates containing 10 mM H_2_DCF-DA. The fluorescence was recorded using Spectra Max M2 multimode microplate reader (Molecular Devices) at 485 nm excitation and 530 nm emission. The change in fluorescence was recorded for 120 min at 20 min intervals at 37 °C. The experiment was performed 3 times independently. To perform the H_2_DCF-DA assay using an image, we transferred 30-40 pretreated *spg-7* and *isp-1* RNAi animals in 500 µl M9 buffer and washed 3 times. Afterward, animals were incubated in 500 µl M9 buffer containing 10 mM H_2_DCF-DA for 1 hour. Animals were washed with M9 buffer at least 3 times, transferred to 2% agarose pads on glass slides, and visualized using a GFP filter and imaged using a 40x objective (Andor, DSD2 spinning disk confocal, Andor, Zyra4.2 camera). To evaluate mitochondrial membrane potential and mass, we used MitoTracker CMXRos (Invitrogen, M7512) and FM green dye (Invitrogen, M7514), respectively, as described in previous studies ^127, 128^. The lyophilized dyes were dissolved in DMSO and made to the final concentration of 100 μM. Animals were pretreated from the L1 stage to *spg-7* and *isp-1* gRNAi. L4 stage animals (n=30-40) were transferred to *spg-7* and *isp-1* RNAi plates with MitoTracker CMXRos or MitoTracker FM green dye and incubated for 48 hours at room temperature under dark conditions. Next, stained animals were transferred to fresh NGM plates (without the dye) for 1 hour to remove the bacterial fluorescent background inside the gut. The animals were observed using a 10x magnification to visualize the whole-body staining and a 40x objective to observe the anterior body, including the intestine.

### ATP assay

We used previously reported protocols with a minor modification ^52^. Approximately 150 age-synchronized animals treated with *spg-7* and *isp-1* RNAi from the L1 stage were prepared.1-day-old adult animals were washed 5 times with M9 buffer to remove the intestinal bacteria and washed with TE buffer. The animals were frozen at −80 °C. The sample was sonicated for 15 seconds with a 15-second interval, followed by boiling for 10 minutes to release ATP and block ATPase activity. Debris was removed by centrifuging at 4 °C for 10 minutes. The supernatant was collected, and the ATP levels were measured using the bioluminescence detection kit (Promega ENLITEN ATP Assay System, FF2000) and Spectra Max M2 multimode microplate reader (Molecular Devices). Luminescence was normalized to protein content measured with a Pierce BCA protein determination kit (Thermo Scientific, 23227).

### mtDNA quantification

mtDNA quantification was performed using a quantitative PCR (qPCR) based method ^129^. About 30 age-synchronized L4 staged *spg-7(gRNAi)* and *isp-1(gRNAi)* animals were collected in 30 µl of lysis buffer (freshly added proteinase K) and frozen immediately at −80 °C for 10 minutes before lysis at 65 °C for 1 hour, followed by 95°C for 15 minutes, and then maintained at 4 °C. Relative quantification was used for determining the fold changes in mtDNA. 2 µl of lysate sample was used as template DNA in each triplicate reaction. qPCR was performed using the SYBR green mixture in a CFX96 Touch Real-Time PCR System (Bio-Rad). Primers (mtDNA target-specific primer) for *cox-4* and *nd-1* were used to determine the mtDNA copy number. The *cox-4* forward primer 5’GCCGACTGGAAGAACTTGTC-3’ and reverse primer 5’-GCGGAGATCACCTTCCAGTA-3’. The nd-1 forward primer 5’-AGCGTCATTTATTGGGAAGAAGAC-3’ and reverse primer 5’-AAGCTTGTGCTAATCCCATAAATGT-3’. All qPCR results were performed in triplicates.

### Quantitative reverse transcriptase PCR

RNA isolation and quantitative reverse transcriptase PCR (qRT-PCR) analysis were performed as previously described ^130^. Animals raised on *spg-7* or *isp-1* gRNAi plates were synchronized by egg prep. Then, total RNA was extracted using the TRIzol (Invitrogen, 15596026) method from age-synchronized animals (approximately 1,500 animals at the 3-day-old adult stage). RNA was purified using a Qiagen Rneasy kit, and 2 µg RNA was used for cDNA synthesis (Thermo Scientific, Verso cDNA synthesis kit, AB1453A). qPCR was performed using SYBR Green master mix (Bio-Rad Laboratories). qPCR primers are listed below. Act-1 Forward 5’-GCTGGACGTGATCTTACTGATTACC-3’, act-1 Reverse 5’-GTAGCAGAGCTTCTCCTTGATGTC-3’, daf-16 Forward 5’-CCAACACATTCATCCCAGTG-3’, daf-16 Reverse 5’-GATGGGATAGAGGTAGCATT-3’, sod-3 Forward 5’-CTGATGGACACTATTAAGCG-3’, sod-3 Reverse 5’-AAGTGGGACCATTCCTTCCAA-3’, gst-4 Forward 5’-GCTGAGCCAATCCGTAT-3’, gst-4 Reverse 5’-GTAAAATGGGAAGCTGGC-3’, dlk-1 Forward 5’-TCGACGCTATCTCCGAACTT-3’, dlk-1 Reverse 5’-TGCTTGATCTCGGTCTCCTT-3’, hsp-6 Forward 5’-CGAAAGCTATTTGGGAACCA-3’, hsp-6 Reverse 5’-GCTCGTTGATGACACGAAGA-3’, hsp-60 Forward 5’-CCGTCTCTGTCACTATGGGC-3’, hsp-60 Reverse 5’-CTCGAATCCCTCTTTGGCGA-3’, *polg-1* Forward 5’-GTTACGGCCGACGAGATACG-3’, *polg-1* Reverse 5’-CGTAGCTTCCGGACTCCAAA-3’. All qPCR results were performed at least in triplicates.

### Statistics

Statistics were performed using GraphPad PRISM (version 9 and 10). No statistical method was used to pre-determine sample size. For Figures 5D and 7D, outliers were identified using the ROUT method at Q = 1% and excluded from further analysis. Within experimental groups, animals were randomized for each experimental replicate. The qPCR experiments were analyzed using the delta-delta Ct method ^121^. All experiments were reliably reproduced at least 3 times independently. For the qPCR assays, in cases where the control and subjects were not paired, the average control delta Ct value was utilized to calculate delta-delta Ct values. However, if the control and experimental groups were paired, individual control delta Ct values were used to calculate delta-delta Ct values for each paired experimental group.

---

**Figure S1.**
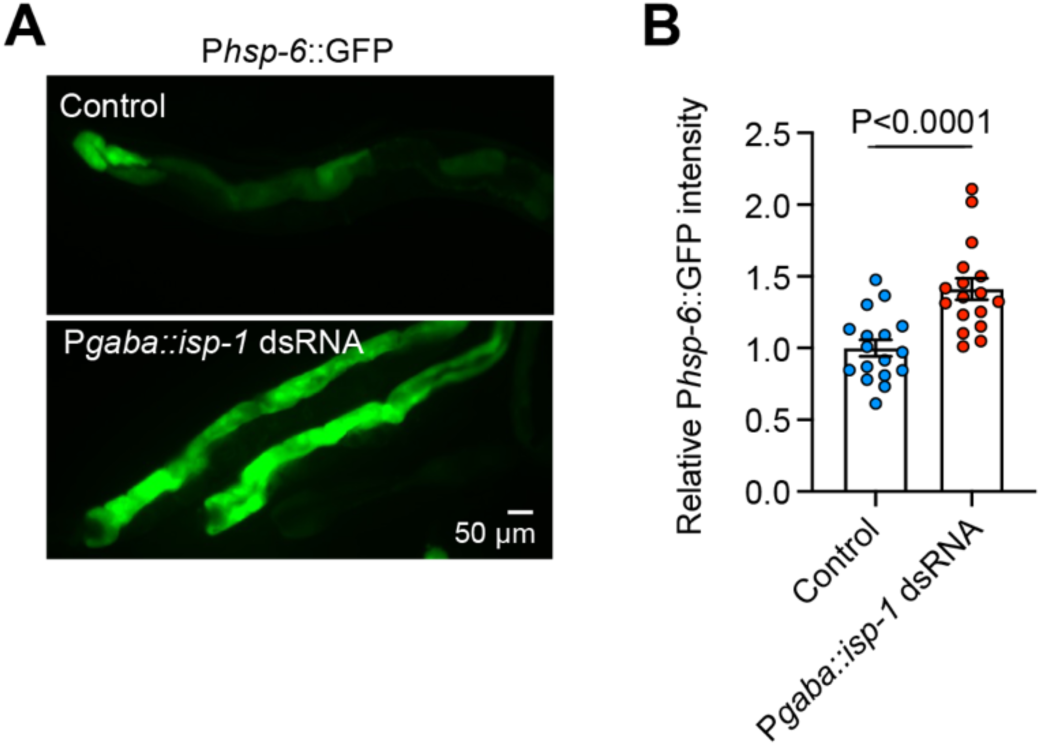
(Related to Figure 1). Global induction of mitoUPR reporter upon *isp-1* dsRNAs expression in GABAergic neurons of wild-type animals. (A) Representative image showing the increased expression of P*hsp-6::gfp* transgene in the animals with transcription of *isp-1* sense and antisense RNAs *in vivo* under the GABAergic neuron-specific promoter. Note that the N2 wild-type background allowed the systemic RNAi effects on peripheral tissues. (B) Quantitation of the P*hsp-6::*GFP fluorescence intensity in the intestine showing the non-cell-autonomous effects of *in vivo* transcription of *isp-1* dsRNAs in GABAergic neurons. Each dot represents an individual animal. Data were expressed as means ± SEM. A two-tailed Mann–Whitney test.

**Figure S2.**
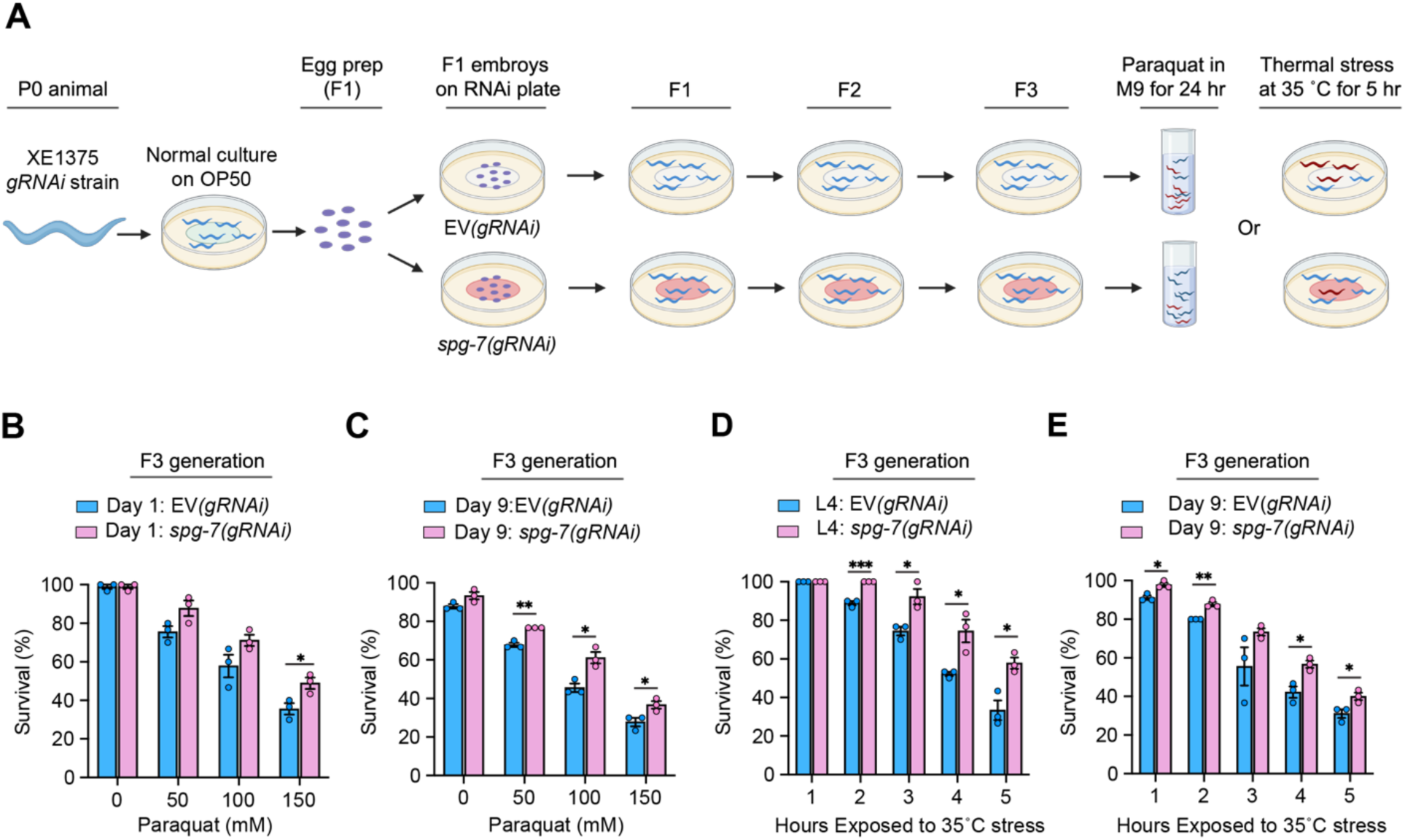
(Related to Figure 2). Continuous gRNAi against *spg-7* over three generations led to altered organismal stress resistance. (A) A diagram showing the experimental procedure of paraquat and thermal stress response assays in *spg-7(gRNAi)* animals after continuous RNAi for three generations. (B and C) Quantification of paraquat stress assays in *spg-7(gRNAi)* animals at 1-day or 9-day-old adult stages. (D and E) Quantification of thermal stress assays in *spg-7(gRNAi)* animals at the L4 and 9-day-old adult stages. Animals were exposed to 35 °C for 5 hours. Data were expressed as means ± SEM. *P.<0.05, **P.<0.005, ***P.<0.005, ****P.<0.001; two-tailed student’s t-test. Created with https://www.biorender.com/.

**Figure S3.**
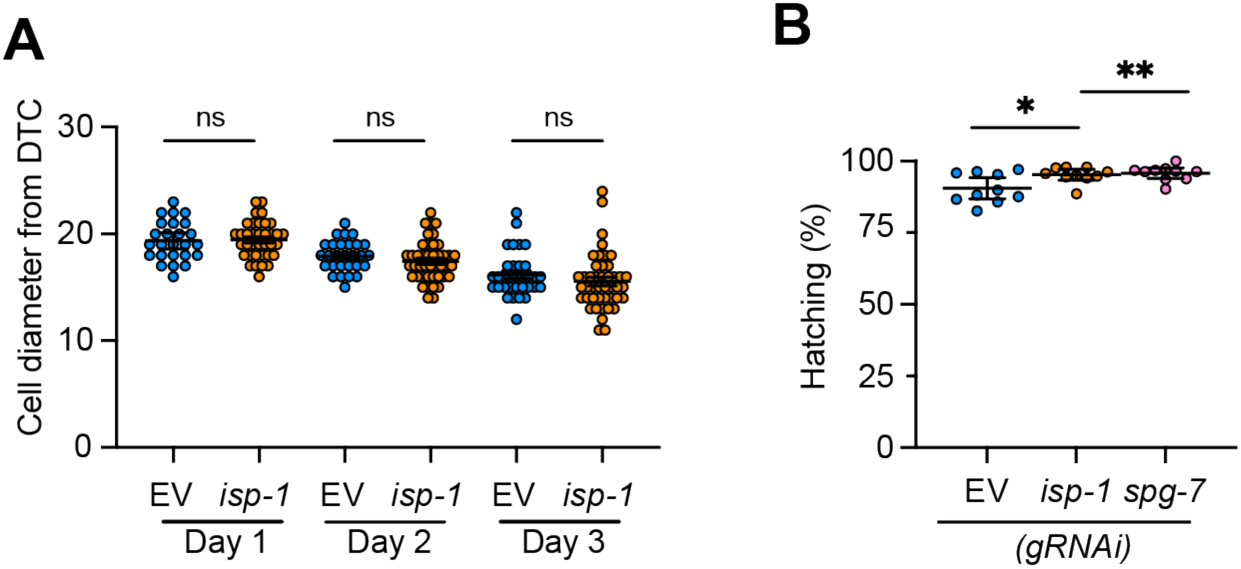
(Related to Figure 3). Non-cell autonomous effects of GABA neuronal mitochondrial stress on the reproductive system. (A) Diameter of the mitotic area in the distal gonad in *isp-1(gRNAi)* animals. (B) Hatching rates of embryos in *isp-1(GABA-RNAi)* and *spg-7(GABA-RNAi)* animals. *P.<0.05, **P.<0.005; a two-tailed Mann–Whitney test for A; ne-way ANOVA test for B. Data were expressed as means ± SEM.

**Figure S4.**
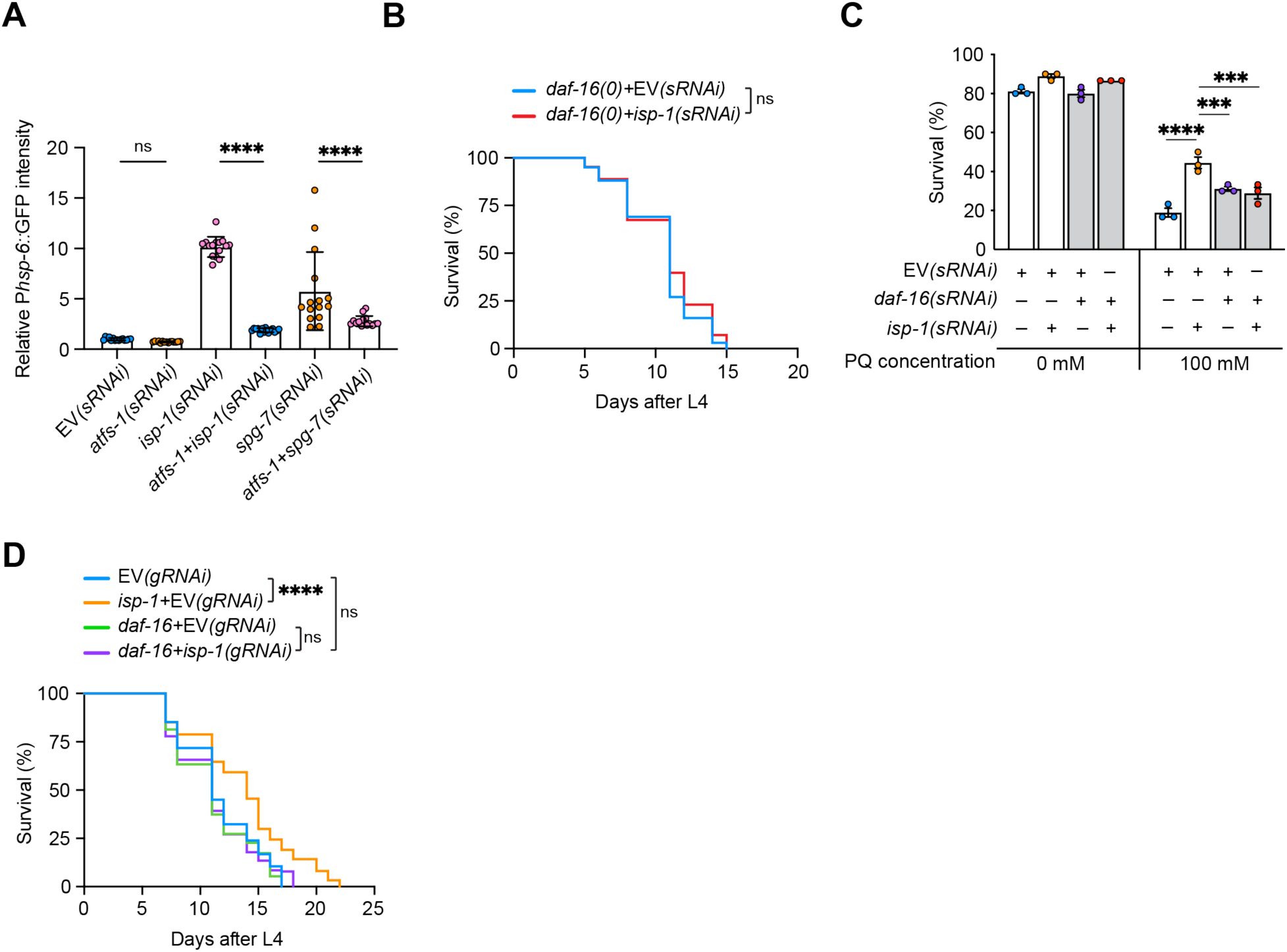
(Related to Figure 6). The role of DAF-16 in the non-cell autonomous effects of GABA neuronal mitochondrial stress. (A) Quantification of the P*hsp-6*::GFP intensity in the indicated sRNAi strains. (B and C) Lifespan (B) and paraquat tolerance (C) assays of isp-1 sRNAi under the *daf-16(mgDf47)* null mutant background. (D) Lifespan was tested after gRNAi induction by feeding GABAergic neuronal-specific RNAi strain (XE1375) worms with a 50:50 mixture of bacteria producing either EV, *isp-1*, or *daf-16* dsRNAs.*P.<0.05, **P.<0.005, ***P.<0.005, ****P.<0.001; a two-tailed Mann–Whitney test was used for A; Log-rank (Mantel-Cox) test for B and D; one-way ANOVA test for C. Data were expressed as means ± SEM.

**Figure S5.**
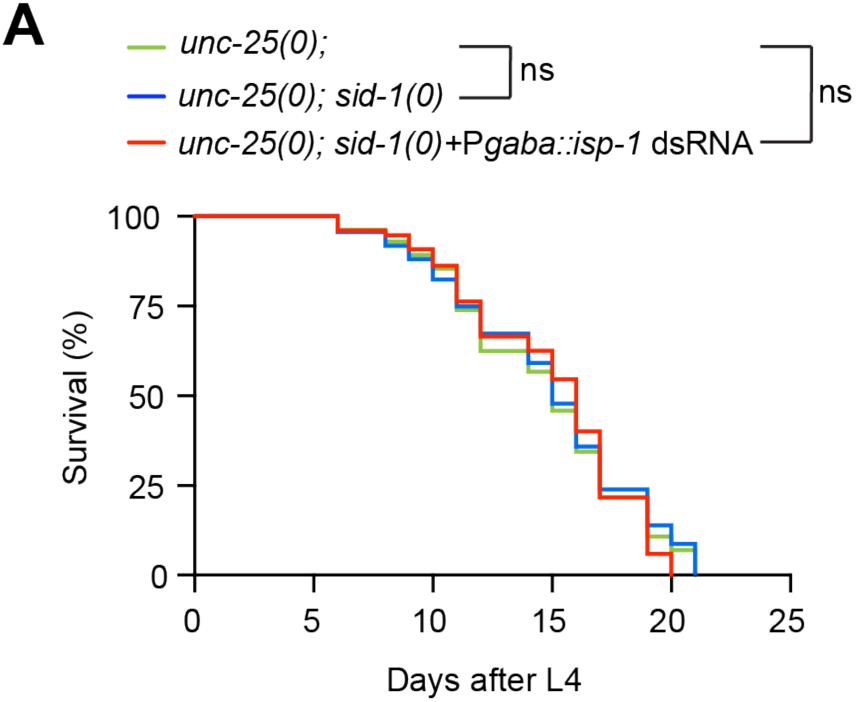
(Related to Figure 7): Assessing the impact of mitochondrial stress in GABAergic neurons on lifespan in *unc-25* null mutants. (A) Repeated lifespan analysis of *unc-25(qt9)* mutants with and without *isp-1(sRNAi)*. The significance was analyzed by a Log-rank (Mantel-Cox) test.

**Figure S6.**
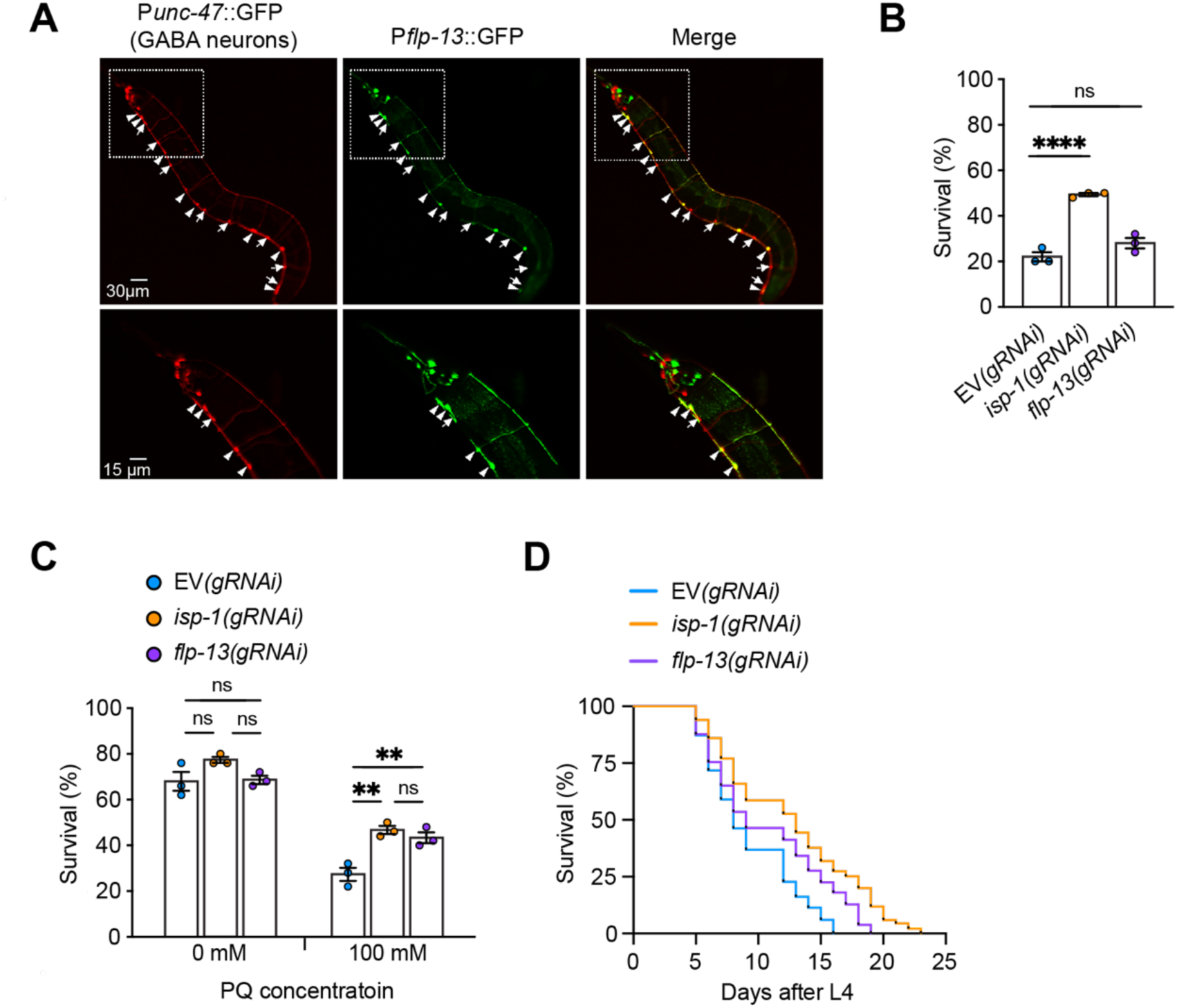
(Related to Figure 8). The effects of single GABA-RNAi inhibition against *flp-13* **and *isp-1* on lifespan and stress response.** (A) Representative images of P*flp-13*::GFP expression. GABA neurons are visualized by mCherry derived by *unc-47* promoter. Arrowheads indicate the cell body of GABA neurons on the ventral nerve cord with GFP expression. Arrows indicate the cell bodies only with mCherry signals. The dashed boxes are magnified below. (A, B, and C) Lifespan and stress tolerance were tested after gRNAi induction by feeding GABAergic neuronal-specific RNAi strain (XE1375) worms with bacteria producing either *isp-1* or *flp-13* dsRNAs alone for each condition. (B) The survival rate after incubation at 35°C for 5 hours was measured in animals at the 3-day-old adult stage. At least 30 animals were tested for each set. (C) The survival rate of animals cultured in indicated concentrations of paraquat for 24 hours was measured at the 3-day-old adult stage. (D) Lifespan assay of *isp-1(gRNAi)* and *flp-13(gRNAi)* animals. **P < 0.005, ****P < 0.001; one-way ANOVA test for B and C; a Log-rank (Mantel-Cox) test for D. Data were expressed as means ± SEM.

**Table S1.** Related to Figs. 1, 6, 7, S4, and S5. Lifespan summary. For P values, stars in the ‘Genotype’ column are provided to denote the genotype (within the group delimited by color) against which the comparison was performed. Actual P values are provided in the ‘significance’ column. The results indicate the total lifespan summary of triplicates.

| Related Figure | Background genotype | RNAi treatment against | Median lifespan (days) | No. of total worms | P (Log-rank test) |
| --- | --- | --- | --- | --- | --- |
| Figure 1A | Wild-type N2 | EV | 16 | 174 |  |
| | | *isp-1* | 23 | 172 | $P < 0.0001$ |
| Figure 1B | Wild-type N2 | EV | 14 | 150 |  |
| | | *spg-7* | 15 | 153 | $P = 0.0001$ |
| Figure 1D | *wpls36 I; wpSi1 II; eri-1(mg366) IV; rde-1(ne219) V; lin-15B(n744) X* | EV | 9 | 127 |  |
| | | *isp-1* | 14 | 140 | $P < 0.0001$ |
| Figure 1E | *wpls36 I; wpSi1 II; eri-1(mg366) IV; rde-1(ne219) V; lin-15B(n744) X* | EV | 12 | 72 |  |
| | | *spg-7* | 16 | 69 | $P < 0.0001$ |
| Figure 1G | *Wild-type N2 |  | 15 | 144 |  |
| | ** *sid-1(qt9)* | | 14.5 | 140 | *$P = 0.59$ |
| | *sid-1(qt9)* | *Pgaba::isp-1* dsRNA | 16 | 151 | *$P < 0.0001$<br>**$P < 0.0001$ |
| Figure 6A | *wpls36 I; wpSi1 II; eri-1(mg366) IV; rde-1(ne219) V; lin-15B(n744) X* | *EV | 11 | 137 |  |
| | | ** *isp-1* | 12 | 113 | *$P < 0.0001$ |
| | *daf-16(mgdf47) wpls36 I; wpSi1 II; eri-1(mg366) IV; rde-1(ne219) V; lin-15B(n744) X* | ***EV | 11 | 117 | *$P = 0.77$ |
| | | *isp-1* | 11 | 148 | *$P = 0.045$<br>**$P < 0.0001$<br>***$P = 0.097$ |
| Figure 7A | *unc-25(e156); sid-1(qt9)* |  | 19.50 | 96 |  |
| | *unc-25(e156); sid-1(qt9)* | *Pgaba::isp-1* dsRNA | 20 | 138 | $P=0.0029$ |
| Figure 8A | *Wild-type N2 |  | 14 | 126 |  |
| | ** *unc-31(e928)* | | 17 | 145 | *$P < 0.0001$ |
| | *** *sid-1(qt9)* | *Pgaba::isp-1* dsRNA | 16 | 154 | *$P < 0.0001$<br>**$P = 0.11$ |
| | *unc-31(e928); sid-1(qt9)* | *Pgaba::isp-1* dsRNA | 16 | 151 | *$P < 0.0001$<br>**$P = 0.04$<br>***$P = 0.59$ |
| Figure 8B |  | *EV | 8 | 149 |  |
|  | *wpls36 I; wpSi1 II; eri-1(mg366) IV; rde-1(ne219) V; lin-15B(n744) X* | *\*\*isp-1+EV* | 9 | 149 | *\*P < 0.0001* |
|  |  | *\*\*\*flp-13+EV* | 13 | 152 | *\*P < 0.0001*<br>*\*\*P = 0.008* |
|  |  | *flp-13+isp-1* | 14 | 135 | *\*P < 0.0001*<br>*\*\*P < 0.0001*<br>*\*\*\*P = 0.02* |
| Figure S4B | *daf-16(mgdf47)* | *\*EV* | 11 | 100 |  |
|  | *daf-16(mgdf47)* | *isp-1* | 11 | 126 | *\*P=0.16* |
| Figure S4D | *wpls36 I; wpSi1 II; eri-1(mg366) IV; rde-1(ne219) V; lin-15B(n744) X* | *\*EV* | 11 | 142 |  |
|  |  | *\*\*isp-1+EV* | 14 | 147 | *\*P = 0.0001* |
|  |  | *\*\*\*daf-16+ EV* | 11 | 150 |  |
|  |  | *daf-16 + isp-1* | 11 | 140 | *\*P = 0.76*<br>*\*\*P < 0.0001*<br>*\*\*\*P = 0.48* |
| Figure S5A | *\*unc-25(e156)* |  | 15 | 157 |  |
|  | *\*\*unc-25(e156); sid-1(qt9)* |  | 16 | 152 | *\*P=0.72* |
|  | *unc-25(e156); sid-1(qt9)* | *Pgaba::isp-1 dsRNA* | 15 | 159 | *\*P=0.56*<br>*\*\*P=0.31* |
| Figure S6D | *wpls36 I; wpSi1 II; eri-1(mg366) IV; rde-1(ne219) V; lin-15B(n744) X* | *\*EV* | 8 | 149 |  |
|  |  | *\*\*isp-1* | 13 | 135 | *\*P<0.0001* |
|  |  | *flp-13* | 9 | 155 | *\*P<0.0001*<br>*\*\*P=0.0001* |

